# Chromosome 1q amplification perturbs a ceRNA network to promote melanoma metastasis

**DOI:** 10.1101/2021.11.14.468531

**Authors:** Xiaonan Xu, Kaizhen Wang, Olga Vera, Akanksha Verma, Olivier Elemento, Xiaoqing Yu, Florian A. Karreth

**Affiliations:** Department of Molecular Oncology, H. Lee Moffitt Cancer Center and Research Institute, Tampa, Florida 33612, USA; Caryl and Israel Englander Institute for Precision Medicine, Institute for Computational Biomedicine, Department of Physiology and Biophysics, Weill Cornell Medicine, New York, NY, 10065, USA; Department of Biostatistics and Bioinformatics, H. Lee Moffitt Cancer Center and Research Institute, Tampa, Florida 33612, USA

**Keywords:** ceRNA, miRNA, melanoma, metastasis, CEP170, NUCKS1, ZC3H11A, chromosome 1q amplification

## Abstract

Somatic copy number alterations (CNAs) promote cancer, but the underlying driver genes are often not obvious when only the functions of the encoded proteins are considered. mRNAs can act as competitive endogenous miRNA sponges (ceRNAs) to post-transcriptionally regulate gene expression in a protein coding-independent manner. However, whether ceRNAs contribute to the oncogenic effects of CNAs is unknown. We report that chromosome 1q gains promote melanoma progression and metastasis at least in part through overexpression of three mRNAs with ceRNA activity: *CEP170, NUCKS1*, and *ZC3H11A*. Genetic studies reveal that these ceRNAs enhance melanoma metastasis by sequestering tumor suppressor miRNAs, thereby alleviating the repression of several pro-metastatic target genes. This regulatory RNA network is evident in other cancer types, suggesting that chromosome 1q ceRNA deregulation is a common driver of cancer progression. Taken together, our work demonstrates that ceRNAs mediate the oncogenicity of somatic CNAs.

## INTRODUCTION

Malignant melanoma remains the leading cause of skin cancer-related deaths, almost invariably due to metastases to vital organs. Accordingly, significant efforts have focused on understanding drivers of melanoma progression and metastasis. While genetic events affecting oncogenes and tumor suppressors such as BRAF, NRAS, CDKN2A, and PTEN are well-documented drivers of melanoma development (Cancer Genome Atlas Network, 2015), surprisingly few metastasis-specific mutations have been discovered (Shain et al., 2018; Vergara et al., 2021). Indeed, mutations and pathways thus far implicated in melanoma metastasis are often those that also promote the early stages of the disease (Damsky et al., 2014; Orgaz and Sanz-Moreno, 2013; Turner et al., 2018), as seen in investigations of autochthonous melanoma models (Ackermann et al., 2005; Cho et al., 2015; Damsky et al., 2011; Dankort et al., 2009; Otsuka et al., 1998; Zingg et al., 2018). Interestingly, copy number alterations (CNA) increase with melanoma progression (Shain et al., 2018; Vergara et al., 2021), suggesting copy number gains and losses as potential drivers of melanoma metastasis. CNA may be focal but can involve entire chromosome arms, obscuring the identity of bona fide metastasis driver genes (Beroukhim et al., 2010). In addition, driver genes are classified based on the function of the encoded protein while potential coding-independent functions of their mRNAs are not usually considered as factors contributing to tumorigenesis.

In addition to being blueprints for translation, mRNAs can act as competitive endogenous RNAs (ceRNAs) that function as natural microRNA (miRNA) sponges. Specifically, ceRNAs competitively bind to miRNAs to impair their repressive activity towards their targets (i.e., ceRNA effectors) (Karreth and Pandolfi, 2013; Salmena et al., 2011). Originally described for pseudogenes (Karreth et al., 2015; Poliseno et al., 2010), ceRNA activity has also been observed for dozens of long noncoding RNAs (lncRNAs) and circular RNAs (circRNAs). Several mRNAs have been reported to promote cancer as ceRNAs, including *MYCN* (Powers et al., 2016), *TYRP1* (Gilot et al., 2017), and several mRNAs that control the expression of PTEN (Karreth et al., 2011; Sumazin et al., 2011; Tay et al., 2011). These studies support the notion that mRNAs can function as ceRNAs to regulate gene expression in cancer, yet it is unknown if ceRNA deregulation is a common phenomenon underlying the oncogenic effects of CNAs. We tested whether copy number gains promote melanoma metastasis via overexpression of oncogenic ceRNAs, revealing a chromosome 1q-encoded, pro-metastatic ceRNA network that is controlled by the *CEP170, NUCKS1*, and *ZC3H11A* mRNAs. Notably *CEP170, NUCKS1*, and *ZC3H11A* are co-gained in several cancers where their expression correlates with upregulation of pro-metastatic ceRNA effectors, suggesting this potent ceRNA network plays oncogenic roles across cancer types. Our study challenges the notion that somatic CNAs promote cancer predominantly through their encoded proteins and establishes ceRNAs as potent drivers underlying the oncogenicity of somatic CNAs.

## RESULTS

### Copy number gains and overexpression of 1q ceRNA genes are associated with melanoma metastasis

To identify gained/amplified genes with putative ceRNA potential, we analyzed a TCGA Skin Cutaneous Melanoma dataset containing 366 metastatic melanoma cases. Given that gained/amplified genes with high basal expression levels are more likely to elicit prominent ceRNA effects by contributing a large number of miRNA binding sites to their ceRNA networks, we first identified genes as recurrently gained in ≥3% of cases and focused on the top 20% highest expressed genes. This analysis yielded 211 candidate genes (**Table S1**). To define putative ceRNA networks we calculated the correlation of the 211 gained candidate genes with the protein-coding transcriptome (20,291 transcripts). We identified 34,285 significant gene pairs with FDR <0.05 and Spearman correlation ≥0.5. These gene pairs were then tested for enrichment of known miRNA binding sites based on TargetScan algorithm predictions (release 7.2) (Agarwal et al., 2015). To capture highly conserved binding sites for the miRNA families (i.e. miRNA Response Elements, MRE), we used the aggregate P_CT_ score (Friedman et al., 2009) with cut-off of ≥0.2 for every predicted miRNA as a stringent probability metric for conserved targeting (**Table S2**). Out of 34,285 gene pairs, 2,515 gene pairs share at least 4 MREs in their 3’UTRs (**Table S3**) and could thus constitute a complex ceRNA network encompassing 40 putative ceRNAs and 971 effector transcripts (**Figure S1A**). The number of effectors for each ceRNA ranged from 1-454 transcripts, with 9 ceRNAs engaging over 100 effectors (**Figure 1A**).

**Figure 1.**
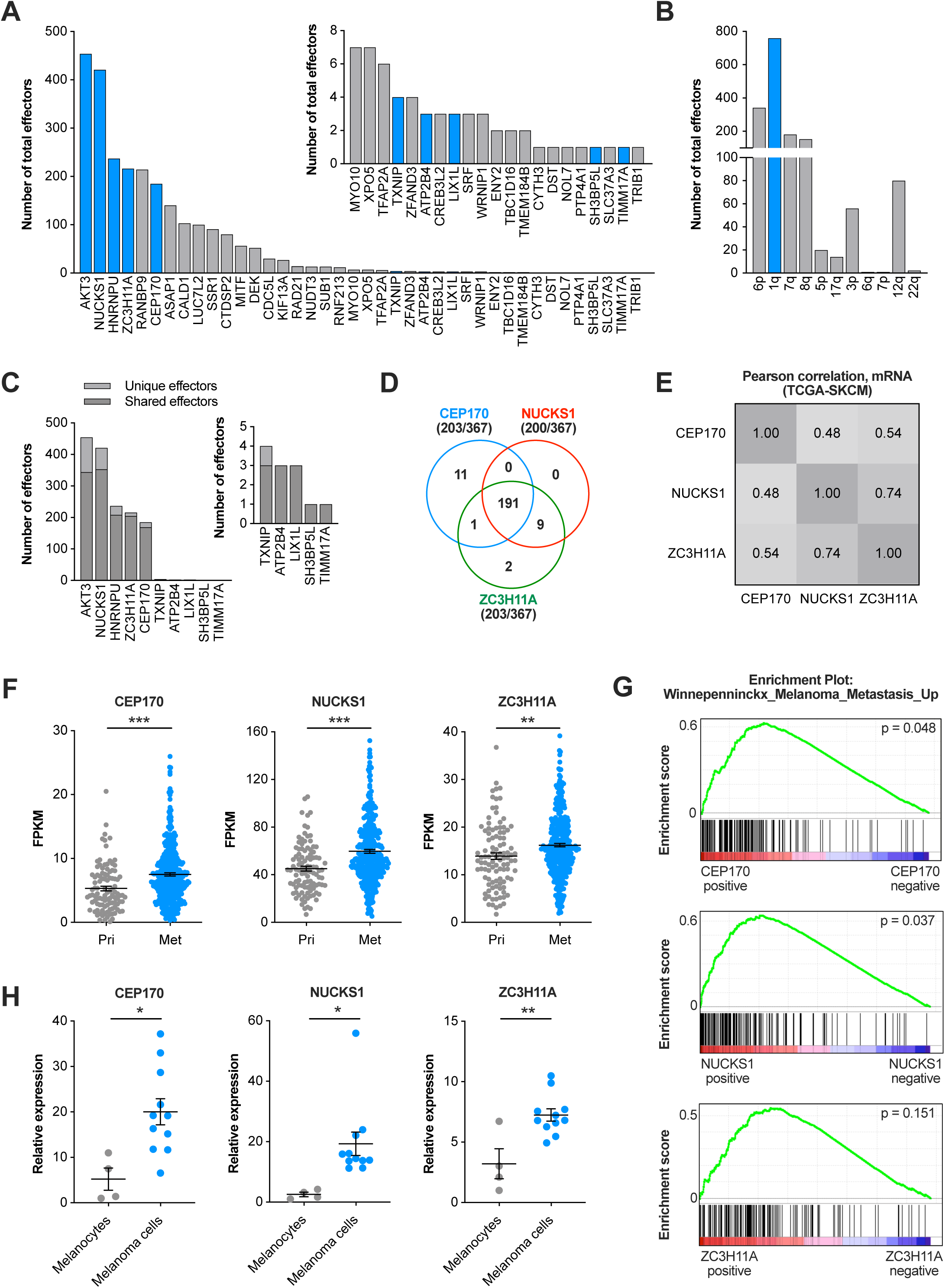
Copy number gains and overexpression of predicted 1q ceRNA genes is associated with melanoma metastasis. (A) The number of total predicted effectors for each ceRNA gene. The insert graph shows a close-up of the 21 bottom-ranked ceRNA genes. Predicted ceRNAs encoded by genes localized on chromosome 1q are highlighted in blue. (B) The number of putative effectors graphed by the genomic localization of their corresponding ceRNA genes. (C) Number of unique and shared effectors of the ceRNAs localized on chromosome 1q. (D) Copy number alterations of *CEP170, NUCKS1*, and *ZC3H11A* in TCGA cutaneous melanoma samples. *CEP170* and *ZC3H11A* are gained/amplified in 203 out of 367 samples, while *NUCKS1* is gained/amplified in 200 samples. All three genes are co-gained/amplified in 191 samples. (E) Correlation of *CEP170, NUCKS1*, and *ZC3H11A* expression in TCGA cutaneous melanoma samples. (F) Expression of *CEP170, NUCKS1*, and *ZC3H11A* in primary and metastatic cutaneous melanoma samples from TCGA. (G) GSEA analysis of TCGA cutaneous melanoma samples associates *CEP170, NUCKS1*, and *ZC3H11A* expression with a melanoma metastasis gene signature. (H) Relative *CEP170, NUCKS1*, and *ZC3H11A* mRNA levels in human melanocyte (Hermes1, Hermes2, Hermes3A, Hermes4B) and melanoma (WM35, WM115, WM164, WM266.4, 451Lu, 501Mel, SbCl2, SKMel28, A375, WM793, 1205Lu) cell lines. In (F, H), mean ± SEM is shown; * p < 0.05, ** p < 0.01, *** p < 0.001. See also Figure S1.

To further systematically prioritize candidate ceRNAs, we evaluated the connectivity of ceRNA nodes in the network using the Hyperlink-Induced Topic Search (HITS) algorithm (**Figure S1B**), which revealed the 5 of the top 6 highly ranked putative ceRNA genes located on chromosome 1q (**Table S4**). The 40 predicted ceRNA genes localize to only 11 genomic regions, with chromosome 6p and 1q harboring 13 and 10 putative ceRNA genes, respectively (**Figure S1C**). This raised the possibility of co-gained ceRNA genes coordinately regulating the same effectors. ceRNA genes located within a genomic region connect to similar effectors, with the greatest number of shared effectors observed for chromosome 1q ceRNAs (**Figure S1D**). Among ceRNA genes localized in the chromosome 1q amplicon (1q^AMP^ ceRNAs), *AKT3, NUCKS1, HNRNPU, ZC3H11A*, and *CEP170* showed the greatest number of total and shared effectors (**Figure 1C**).

This ceRNA prediction was based on annotated full-length 3’UTRs. However, alternative polyadenylation (APA) can shorten 3’UTRs and thereby limit their ability to sequester miRNAs. Using data from the APAatlas (https://hanlab.uth.edu/apa/) (Hong et al., 2019), we assessed the prevalence of APA across several tissue types for *AKT3, NUCKS1, HNRNPU, ZC3H11A*, and *CEP170*. This analysis revealed almost universal APA for *HNRNPU*, a wide range of polyA site usage for *NUCKS1*, and predominant usage of canonical polyA sites for *AKT3, ZC3H11A*, and *CEP170* (**Figure S1E**). Given shortened 3’UTRs for *HNRNPU* transcripts, this mRNA was excluded from further analyses. Moreover, there were no significant biological effects of ectopic expression of the 3’UTR or coding sequence of *AKT3* on two human melanoma cell lines (**Figure S1F-H**). Consequently, the 1q^AMP^ ceRNAs *CEP170, NUCKS1*, and *ZC3H11A* were assessed for their roles in melanoma.

Chromosome 1q copy number gains correlate with melanoma stage and occur in over 25% of metastatic melanoma (Bastian et al., 1998; Fountain et al., 1990; Mertens et al., 1997; Thompson et al., 1995). Accordingly, analysis of a TCGA Skin Cutaneous Melanoma dataset revealed the 1q^AMP^ ceRNA genes *CEP170, NUCKS1*, and *ZC3H11A* show co-occurrence of copy number gains or amplification in approximately 52% (191 out of 367) of metastatic melanoma cases (**Figure 1D**). Further, copy number gains of chromosome 1q are associated with increased expression of *CEP170, NUCKS1*, and *ZC3H11A* (**Figure S1I**) and the expression levels of these 3 genes are significantly correlated (**Figure 1E**). Expression of *CEP170, NUCKS1*, and *ZC3H11A* is also significantly increased in metastases compared to primary melanomas (**Figure 1F**). In accord with these findings, *CEP170* and *NUCKS1* expression is associated with a gene set enrichment analysis metastasis signature, and *ZC3H11A* also exhibited a trend towards association with the metastasis signature (**Figure 1G**). Moreover, expression of *CEP170, NUCKS1*, and *ZC3H11A* is increased in human melanoma cells compared to human immortalized melanocyte cell lines (**Figure 1H**, **Table S5**). Therefore, our data indicate that copy number gains and overexpression of the putative 1q^AMP^ ceRNAs *CEP170, NUCKS1*, and *ZC3H11A* are associated with melanoma progression and metastasis.

### 1q^AMP^ ceRNAs have oncogenic effects in melanoma cells in vitro

To assess the functional potential of *CEP170, NUCKS1*, and *ZC3H11A* as ceRNAs, we tested if forced overexpression of their 3’UTRs impacts the phenotype of human melanoma cells. We generated lentiviral vectors that express the 3’UTRs of *CEP170, NUCKS1*, and *ZC3H11A* linked to a GFP cDNA to prevent nonsense-mediated decay, ensure proper folding of the 3’UTRs, and enable tracing of cells expressing the 3’UTRs. Ectopic expression of these constructs in the human melanoma cell lines A375 and WM793 that lack chromosome 1q CNA resulted in 10-40-fold overexpression of the *CEP170, NUCKS1*, and *ZC3H11A* 3’UTRs (**Figure S2A**). While overexpression of the *CEP170, NUCKS1*, or *ZC3H11A* 3’UTRs had only modest effects on melanoma cell proliferation (**Figure S2B and S2C**), there were marked increases in colony number and size under anchorage-independent conditions in soft agar (**Figure 2A**). Further, overexpression of these 3’UTRs also enhanced melanoma cell migration and invasion in transwell assays (**Figure 2B** **and 2C**). Notably, combined overexpression of the *CEP170, NUCKS1*, and *ZC3H11A* 3’UTRs further increased anchorage-independent growth, migration, and invasion (**Figure 2A-C**), suggesting cooperative effects of these ceRNAs.

**Figure 2.**
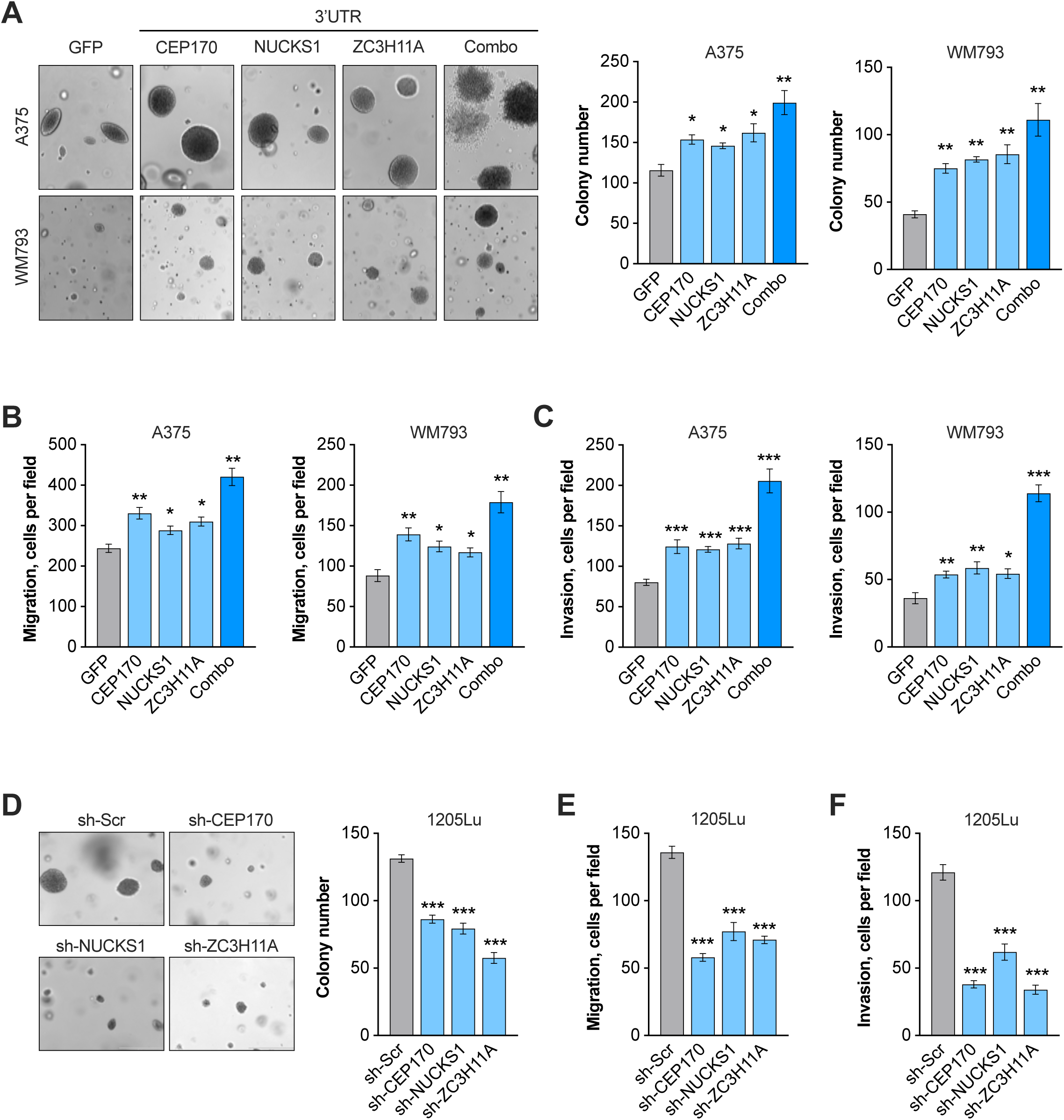
1q^AMP^ ceRNAs have oncogenic effects in melanoma cells in vitro. (A) Anchorage-independent growth of A375 and WM793 cells expressing GFP, GFP-CEP170-3’UTR, GFP-NUCKS1-3’UTR, GFP-ZC3H11A-3’UTR, or the combination of the three 3’UTRs (Combo). Representative images are show on the left, quantifications are shown on the right. (B) Quantification of migrated cells in transwell migration assays using the cell lines shown in (A). (C) Quantification of invasive cells in transwell invasion assays using the cell lines shown in (A). (D) Anchorage-independent growth of 1205Lu cells expressing shRNAs targeting *CEP170, NUCKS1*, or *ZC3H11A*. A scrambled shRNA (sh-Scr) was used as negative control. Representative images are show on the left, quantifications are shown on the right. (E) Quantification of migrated cells in transwell migration assays using the cell lines shown in (D). (F) Quantification of invasive cells in transwell invasion assays using the cell lines shown in (D). N = 3 biological replicates; two-sided t test. Values represent mean ± SEM; * p < 0.05, ** p < 0.01, *** p < 0.001. See also Figure S2.

To affirm the oncogenic potential of 1q^AMP^ ceRNAs, we used shRNAs to silence endogenous *CEP170, NUCKS1*, or *ZC3H11A* expression in metastatic 1205Lu human melanoma cells (**Figure S2D**). Reduced expression of endogenous *CEP170, NUCKS1*, or *ZC3H11A* impaired anchorage-independent growth in soft agar, migration and invasion of 1205Lu cells (**Figure 2D-F**). To test if these effects were mediated by the encoded CEP170, NUCKS1, and ZC3H11A proteins, we overexpressed the coding sequences (CDS) of the 1q^AMP^ ceRNA genes in A375 and WM793 cells. Interestingly, the CEP170, NUCKS1, and ZC3H11A CDS had no effect on growth in soft agar or on transwell migration and invasion (**Figure S2E and 2F**), demonstrating that *CEP170, NUCKS1*, and *ZC3H11A* augment aggressive melanoma cell phenotypes in vitro in a 3’UTR-dependent but protein coding-independent manner.

### 1q^AMP^ ceRNAs augment melanoma metastasis in vivo

To test the role of the 3’UTRs of *CEP170, NUCKS1*, and *ZC3H11A* in promoting melanoma metastasis, A375 and WM793 melanoma cells harboring luciferase were engineered to express the ectopic *CEP170, NUCKS1*, and *ZC3H11A* 3’UTRs. A375 cells were subcutaneously injected into the flanks of NSG mice for spontaneous metastasis assays, while WM793 cells were intravenously inoculated into the tail veins of NSG mice. Notably, bioluminescence imaging revealed that overexpression of the *CEP170, NUCKS1*, and *ZC3H11A* 3’UTRs significantly increased tumor burden in the lungs in the A375 and WM793 metastasis models (**Figures 3A** **and 3B**). H&E staining corroborated the increased lung metastasis burden upon 1q^AMP^ ceRNA overexpression (**Figure 3C** **and 3D**). While combined overexpression of the *CEP170, NUCKS1*, and *ZC3H11A* 3’UTRs also enhanced lung metastasis, the effect was comparable to overexpression of individual 3’UTRs. Finally, silencing of *CEP170, NUCKS1*, or *ZC3H11A* in luciferase-expressing 1205Lu melanoma cells significantly impaired their lung metastatic potential in intravenously inoculated NSG mice (**Figure 3E** **and 3F**). Thus, 1q^AMP^ ceRNAs promote the metastatic program of melanoma cells.

**Figure 3.**
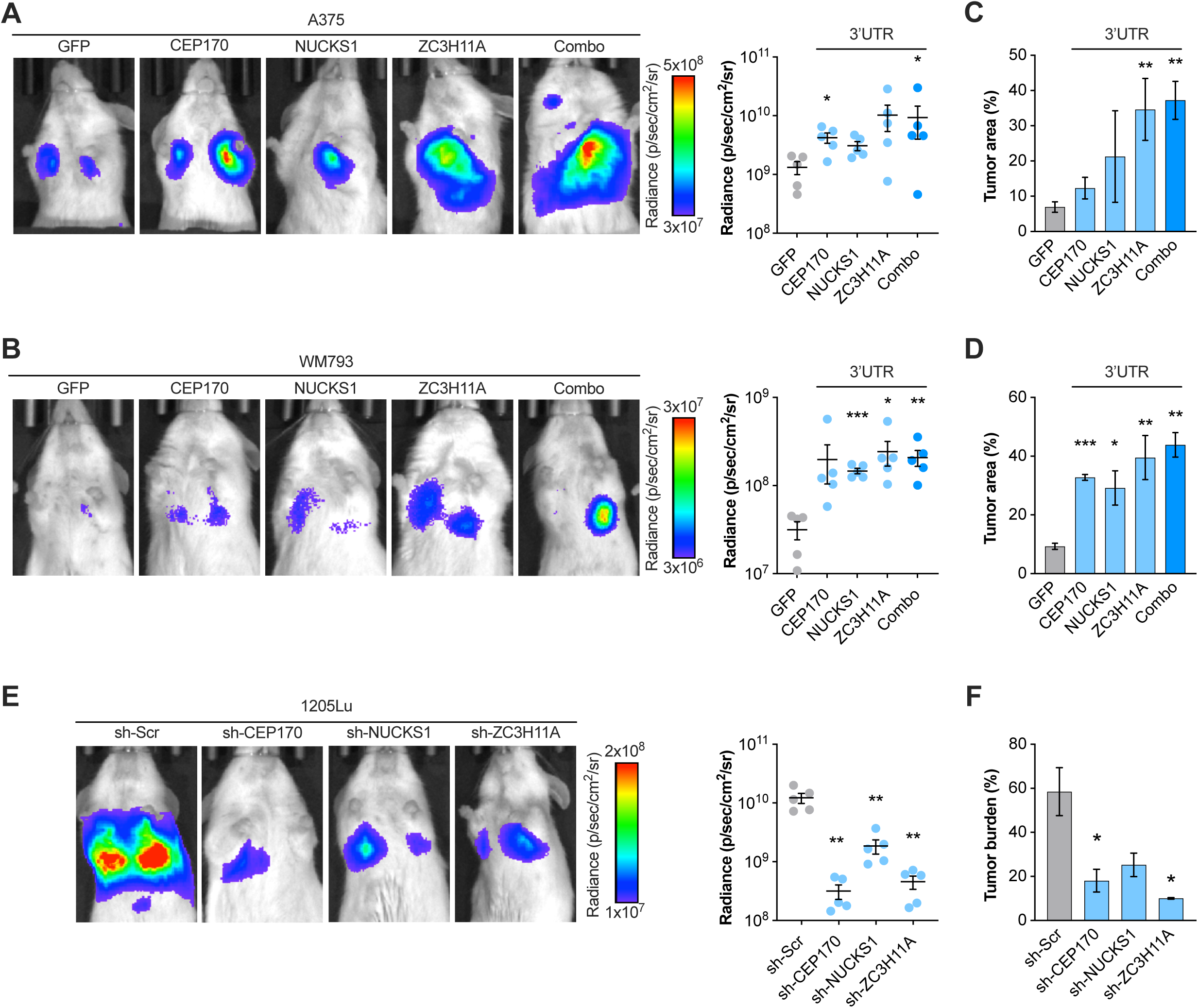
1q^AMP^ ceRNAs enhance melanoma metastasis in vivo. (A-B) Luciferase tagged A375 (A) and WM793 (B) cells expressing GFP, GFP-CEP170-3’UTR, GFP-NUCKS1-3’UTR, GFP-ZC3H11A-3’UTR, or the combination of the three 3’UTRs (Combo) were injected into NSG mice subcutaneously (A375) or intravenously (WM793). Representative bioluminescence images (left) and quantification of the luminescence signals (right) in lungs after 28 days (A375) or 60 days (WM793) are shown. (C-D) Quantification of lung metastasis burden by H&E staining of the mice injected with A375 (C) or WM793 (D) cells shown in (A) and (B), respectively. (D) Quantification of lung metastasis burden by H&E staining of the mice shown in (B). (E) Luciferase tagged 1205Lu cells expressing shRNAs against *CEP170, NUCKS1*, or *ZC3H11A* or a scrambled control (sh-Scr) were intravenously injected into NSG mice and the luminescence signal in lungs was quantified after 25 days. Representative bioluminescence images (left) and quantification of the luminescence signals (right) are shown. (F) Quantification of lung metastasis burden by H&E staining of the mice injected with 1205Lu cells shown in (E). Each data point in (A,B,E) represents one mouse; (C,D,F) N = 3-4 biological replicates. Two-sided t test; values represent mean ± SEM. * p < 0.05, ** p < 0.01, *** p < 0.001.

### 1q^AMP^ ceRNAs sequester tumor suppressive miRNAs

To define the mechanism underlying the oncogenic effect of the 1q^AMP^ ceRNAs, we assessed the ability of the *CEP170, NUCKS1*, and *ZC3H11A* 3’UTRs to function as miRNA sponges. To this end we created a series of luciferase reporters harboring binding sites for the miRNAs predicted to engage in the 1q^AMP^ ceRNA network (**Table S6**). Luc-MRE reporters were expressed in WM793 and A375 melanoma cells with our without the *CEP170, NUCKS1*, or *ZC3H11A* 3’UTRs (**Figure S3A**). This identified miRNAs that are expressed and active (i.e., Luc-MRE reporter activity is reduced compared to empty Luc control reporter) as well as those that are sequestered by the 1q^AMP^ ceRNAs (i.e., Luc-MRE reporter activity is increased by the *CEP170, NUCKS1*, or *ZC3H11A* 3’UTRs compared to GFP). This analysis revealed that: (i) miR-135, miR-141, miR-144, miR-200a, and miR-203 are sponged by the *CEP170* 3’UTR: (ii) miR-135, miR-137, and miR-203 are sponged by the *NUCKS1* 3’UTR; and (iii) miR-101, miR-125, miR-137, miR-144, miR-200bc, and miR-497 are sponged by the *ZC3H11A* 3’UTR (**Figures 4A-C** **and S3B-D**). Of note, miR-135, miR-137, miR-144, and miR-203 are sequestered by two 3’UTRs, and the combined expression of the 1q^AMP^ ceRNA 3’UTRs further enhanced Luciferase activity of these four Luc-MRE reporters (**Figure S3E**).

**Figure 4.**
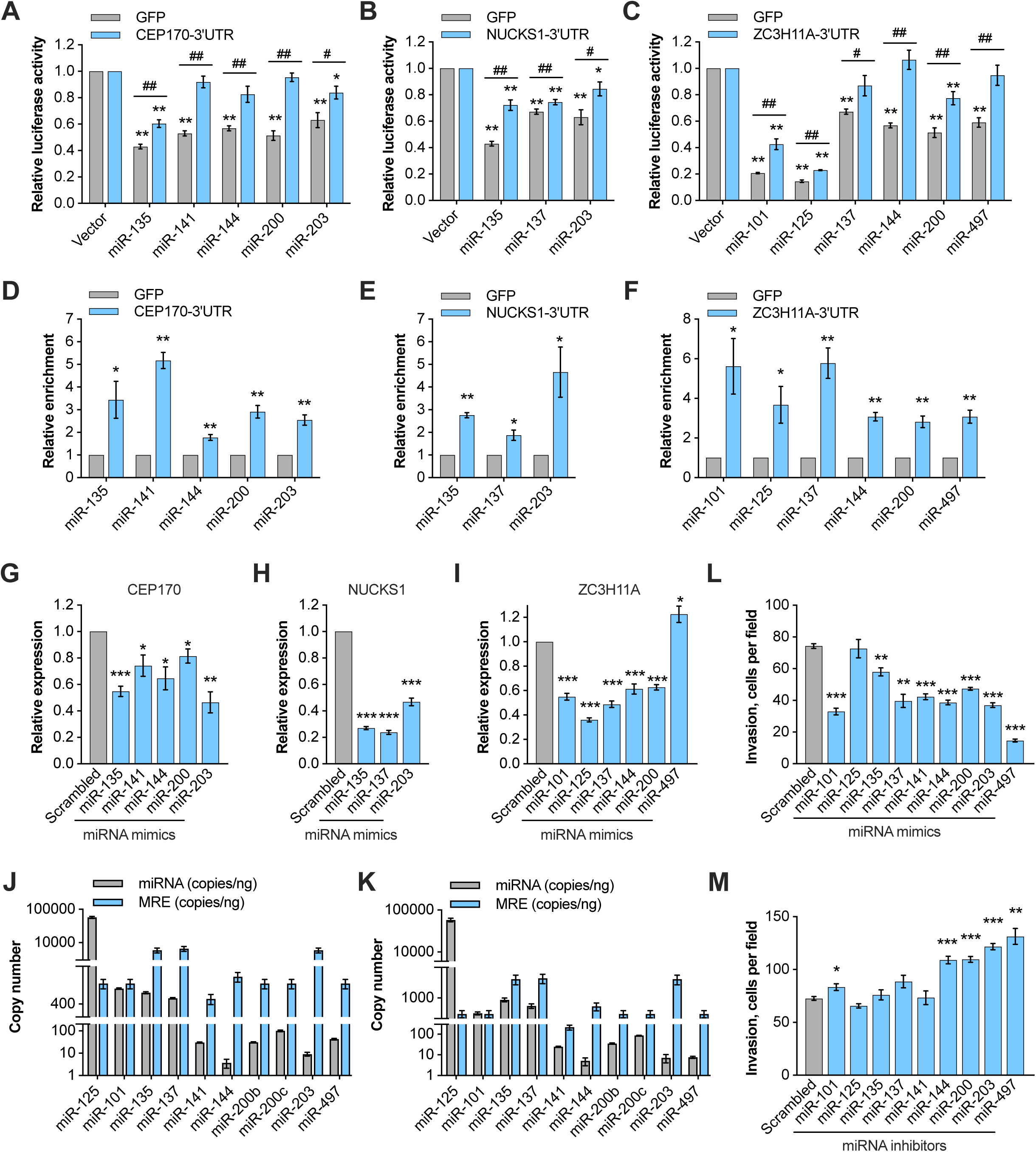
1q^AMP^ ceRNAs sequester tumor suppressor miRNAs. (A) Luciferase assays using dual luciferase reporters harboring MREs for miR-135-5p, miR-141-3p, miR-144-3p, miR-200-3p, or miR-203-3p in WM793 cells expressing GFP or GFP-CEP170-3’UTR. Luciferase activity was normalized to the reporter without MRE (vector). (B) Luciferase assays using dual luciferase reporters harboring MREs for miR-135-5p, miR-137, or miR-203-3p in WM793 cells expressing GFP or GFP-NUCKS1-3’UTR. Luciferase activity was normalized to the reporter without MRE (vector). (C) Luciferase assays using dual luciferase reporters harboring MREs for miR-101-3p, miR-125-5p, miR-137, miR-144-3p, miR-200-3p, or miR-497-5p in WM793 cells expressing GFP or GFP-ZC3H11A-3’UTR. Luciferase activity was normalized to the reporter without MRE (vector). In (A-C), asterisks (*, **) indicate comparison of Luc-MRE reporter vs. empty control Luc reporter (Vector) and pound symbols (^#^, ^##^) indicate comparison of 3’UTR vs. GFP. (D-F) RNA pulldown of GFP-CEP170-3’UTR (D), GFP-NUCKS1-3’UTR (E), or GFP-ZC3H11A-3’UTR (F) followed by RT-qPCR for the indicated miRNAs shows the relative enrichment of bound miRNAs. miRNA levels were normalized to pulldown of the GFP control. (G-I) RT-qPCR showing the relative expression of endogenous *CEP170* (G), *NUCKS1* (H), and *ZC3H11A* (I) in 1205Lu cells upon transfection of the indicated miRNA mimics. (J,K) Copy number of the indicated mature miRNAs and the corresponding MREs present in endogenous *CEP170, NUCKS1, ZC3H11A* in A375 (J) and WM793 (K) cells were quantified by Droplet Digital PCR. (L) Quantification of invasive WM793 cells in transwell invasion assays upon transfection with the indicated miRNA mimics. (M) Quantification of invasive WM793 cells in transwell invasion assays upon transfection with the indicated miRNA inhibitors. N = 3 biological replicates; two-sided t test. Values represent mean ± SEM. * p < 0.05, ** p < 0.01, *** p < 0.001, ^#^ p < 0.05, ^##^ p < 0.01. See also Figure S3.

To test direct interaction of these 9 miRNAs with the 1q^AMP^ ceRNAs, we performed RNA pulldowns using biotinylated probes targeting the GFP sequence in WM793 melanoma cells expressing the *CEP170, NUCKS1*, or *ZC3H11A* 3’UTRs (**Figure S3A**). These analyses revealed that: (i) miR-135, miR-141, miR-144, miR-200a, and miR-203 bound the *CEP170* 3’UTR; (ii) miR-135, miR-137, and miR-203 bound the *NUCKS1* 3’UTR; and (iii) miR-101, miR-125, miR-137, miR-144, miR-200bc, and miR-497 bound the *ZC3H11A* 3’UTR (**Figure 4D-F**). Finally, to further validate the interaction of the 1q^AMP^ ceRNAs with these miRNAs, we transfected miRNA mimics into WM793 melanoma cells. As expected, mimics of: (i) miR-135, miR-141, miR-144, miR-200a, and miR-203 decreased *CEP170* expression; (ii) miR-135, miR-137, and miR-203 mimics decreased *NUCKS1* expression; and (iii) miR-101, miR-125, miR-137, miR-144, miR-200bc, and miR-497 mimics reduced *ZC3H11A* expression (**Figure 4G-I**). Thus, the *CEP170, NUCKS1*, and *ZC3H11A* 3’UTRs directly bind to and selectively impair the activity of 9 miRNAs.

To assess the stoichiometry of 1q^AMP^ ceRNAs and the 9 interacting miRNAs, we determined the absolute copy numbers of the endogenous transcripts in A375 and WM793 melanoma cells by Droplet Digital PCR. Notably, with one exception (miR-125) the *CEP170, NUCKS1*, and *ZC3H11A* mRNAs are equally or more abundant than these target miRNAs (**Figure S3F and S3G**). Moreover, with the exception of miR-125, the number of MREs in the 1q^AMP^ ceRNAs significantly exceeded the number of molecules of their corresponding miRNAs (**Figure 4J** **and 4K**), suggesting that in 1q^AMP^ melanoma abundant *CEP170, NUCKS1*, and *ZC3H11A* mRNAs can competitively sequester miRNAs.

If overexpression of the 1q^AMP^ ceRNAs reduces the activity of the bound miRNAs toward other targets, these miRNAs may have tumor suppressive potential. To test this, we delivered miRNA mimics and inhibitors to WM793 and 1205Lu melanoma cells and assessed the effects on cell invasion. Notably, 8 out of 9 miRNA mimics reduced cell invasion, and 5 (WM793) and 7 (1205Lu) miRNA inhibitors increased cell invasion to varying extents (**Figures 4L, 4M, and S3H**). Finally, the expression of miR-141, miR-200a/b/c, and miR-203a is reduced in metastatic melanoma compared to primary tumors in a TCGA skin cutaneous melanoma dataset, further supporting tumor suppressor roles for these miRNAs (**Figure S3I**). These data indicate that 1q^AMP^ ceRNAs bind to and inhibit miRNAs having tumor suppressor potential in melanoma.

### 1q^AMP^ ceRNAs promote melanoma metastasis in a miRNA-dependent manner

To test if the oncogenic potential of the 1q^AMP^ ceRNAs is dependent on the interaction with miRNAs, we silenced *Dicer* in WM793 melanoma cells to acutely inhibit global miRNA biogenesis (**Figure S4A**). Silencing of *Dicer* abolished the pro-invasion effects of individual and combined 1q^AMP^ ceRNA overexpression (**Figure 5A**). To verify this was through sequestration of the identified tumor suppressor miRNAs, the relevant MREs in the *CEP170, NUCKS1*, and *ZC3H11A* 3’UTR overexpression constructs were mutated to prevent miRNA binding to 3’UTRs. Specifically, we mutated: (i) the miR-135, miR-141/200a, miR-144, and miR-203 MREs in *CEP170*; (ii) the miR-135, miR-137, and miR-203 MREs in *NUCKS1*; and (iii) the miR-101/144, miR-137, miR-200bc, and miR-497 MREs in *ZC3H11A*. While overexpression of the wildtype *CEP170, NUCKS1*, and *ZC3H11A* 3’UTRs increased the invasion of WM793 and A375 melanoma cells, the MRE-mutant 3’UTRs had no effect (**Figures 5B-D** **and S4B-D**). Additionally, the wildtype 3’UTRs but not the MRE-mutant 3’UTRs rescued cell invasion in melanoma cells transfected with the corresponding miRNA mimics (**Figures 5B-D** **and S4B-D**). Expression of wildtype but not MRE-mutant *CEP170, NUCKS1*, or *ZC3H11A* 3’UTR constructs in 1205Lu cells in which the corresponding endogenous 1q^AMP^ ceRNAs were silenced rescued cell invasion in vitro (**Figure 5E**) and lung metastasis following intravenous transplantation in NSG mice (**Figure 5F**). These findings demonstrate that the pro-metastatic potential of the 1q^AMP^ ceRNAs depends on their ability to sequester their respective tumor suppressor miRNA targets.

**Figure 5.**
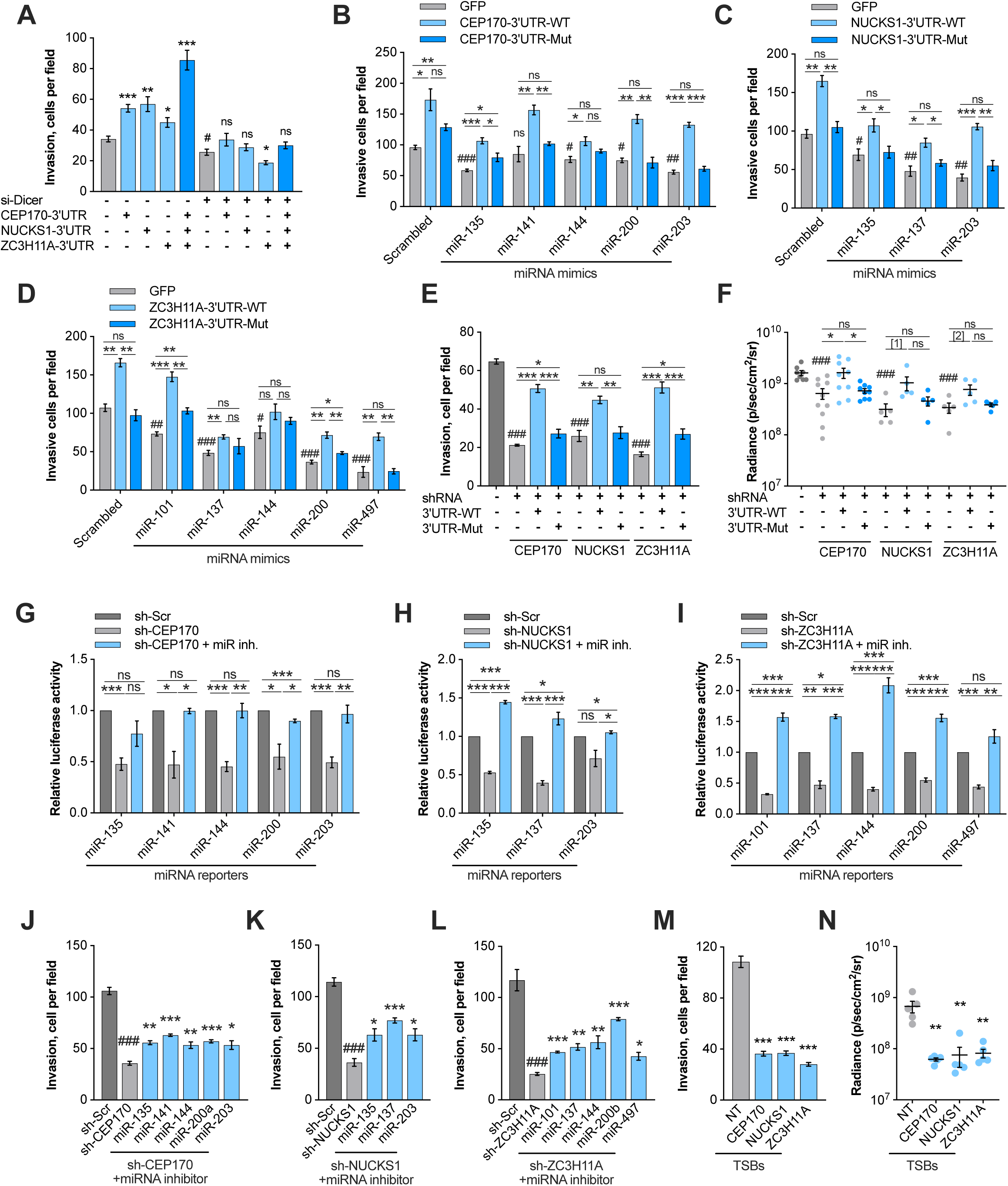
1q^AMP^ ceRNA-directed melanoma cell invasion and metastasis is miRNA dependent. (A) Quantification of transwell invasion of WM793 cells overexpressing GFP (gray bars) or the GFP-linked 1q^AMP^ ceRNA 3’UTRs individually or in combination. WM793 cells were transfected with an siRNA pool targeting Dicer or a non-targeting control siRNA pool. Asterisks indicate comparisons of 3’UTR overexpression (light/dark blue bars) vs. control (gray bars) and pound symbol indicates comparison of si-Dicer vs. non-targeting siRNA in vector control samples (gray bars). (B-D) Quantification of transwell invasion of WM793 cells upon transfection with the indicated miRNA mimics. WM793 cells overexpress GFP, or wildtype or MRE-mutant GFP-CEP170-3’UTR (B), GFP-NUCKS1-3’UTR (C), or GFP-ZC3H11A-3’UTR (D). Asterisks indicate comparisons of GFP vs. 3’UTR-WT vs. 3’UTR-Mut and pound symbols indicate comparisons of miRNA mimics vs. scrambled control in GFP control samples (blue bars). (E) Quantification of transwell invasion of 1205Lu cells in which the 1q^AMP^ ceRNAs were knocked-down followed by restoration with either wildtype or MRE-mutant GFP-CEP170-3’UTR, GFP-NUCKS1-3’UTR, or GFP-ZC3H11A-3’UTR. (F) Luciferase-labeled 1205Lu cells from (E) were intravenously injected into NSG mice and the bioluminescence signal in lungs was quantified after 25 days. In (E-F), asterisks indicate comparisons of control vs. 3’UTR-WT vs. 3’UTR-Mut and pound symbols indicate comparisons of ceRNA shRNAs (light gray bars/dots) vs. control shRNA (dark gray bar/dots). (G-I) Luciferase assays using 1205Lu cells expressing a control shRNA (sh-Scr) or sh-CEP170 (G), sh-NUCKS1 (H), or sh-ZC3H11A (I). The cells were transfected with the corresponding miRNA inhibitors and MRE-Luc reporters. Luciferase activity normalized to the sh-Scr control is shown. (J-L) Quantification of transwell invasion of CEP170- (J), NUCKS1- (K), or ZC3H11A-silenced (L) 1205Lu cells upon transfection with the indicated miRNA inhibitors. Asterisks indicate comparisons of ceRNA shRNA + miRNA inhibitor (blue bars) vs. ceRNA shRNA (light gray bars) and pound symbols indicate comparisons of ceRNA shRNAs (light gray bars) vs. control shRNA (dark gray bars). (M) Quantification of transwell invasion of 1205Lu cells transfected with TSB combinations to block all relevant MREs on endogenous *CEP170, NUCKS1*, or *ZC3H11A*. Asterisks indicate comparisons of TSBs (light blue bars) vs. control TSB (gray bars). (N) Luciferase-labeled 1205Lu cells were transfected with TSB combinations to block MREs on endogenous *CEP170, NUCKS1*, or *ZC3H11A* followed by intravenous injection into NSG mice. Quantification of the luminescence signal in lungs after 25 days is shown. Asterisks indicate comparisons of TSBs (light blue) vs. control TSB (gray). N = 3 biological replicates; two-sided t test. Values represent mean ± SEM. * p < 0.05, ** p < 0.01, *** p < 0.001, ^#^ p < 0.05, ^##^ p < 0.01, ^###^ p < 0.001, ns not significant. See also Figure S4.

The contribution of miR-101, miR-135, miR-137, miR-141, miR-144, miR-200, miR-203, and miR-497 to the oncogenic effects of endogenously expressed 1q^AMP^ ceRNAs was further tested using miRNA inhibitors and Target Site Blockers (TSBs). Knockdown of *CEP170, NUCKS1*, or *ZC3H11A* in 1205Lu melanoma cells reduced the activity of the respective Luc-MRE reporters (**Figure 5G-I**), indicating that silencing of 1q^AMP^ ceRNAs enhances activity of their miRNA targets. Indeed, co-transfection of the corresponding miRNA inhibitors (i.e., [i] miR-135, miR-141, miR-144, miR-200a, or miR-203 inhibitors in *CEP170*-silenced cells; [ii] miR-135, miR-137, or miR-203 inhibitors in *NUCKS1*-silenced cells; and [iii] miR-101, miR-137, miR-144, miR-200bc, or miR-497 inhibitors in *ZC3H11A*-silenced cells) rescued the repression of the Luc-MRE reporters (**Figure 5G-I**). Similarly, the decrease in cell invasion upon 1q^AMP^ ceRNA silencing was partially rescued by the respective miRNA inhibitors (**Figure 5J-L**).

TSBs block endogenous ceRNA-miRNA interactions and thereby reduce ceRNA-mediated miRNA sequestration. Delivery of TSBs to 1205Lu melanoma cells targeting the respective MREs on *CEP170, NUCKS1*, or *ZC3H11A* affirmed the miRNA sponge effects of endogenous 1q^AMP^ ceRNAs, where the individual TSBs inhibited invasion of 1205Lu melanoma cells (**Figure S4H-J**). To test the effect of blocking the binding of all relevant miRNAs to *CEP170, NUCKS1*, or *ZC3H11A*, we delivered combinations of TSBs to 1205Lu cells. Phenocopying the effects of 1q^AMP^ ceRNA silencing, TSBs significantly reduced the invasion of 1205Lu cells in transwell assays (**Figure 5M**) and the seeding to the lungs upon intravenous transplantation into NSG mice (**Figure 5N**). Thus, 1q^AMP^ ceRNAs promote melanoma progression and lung metastasis by sequestering 8 tumor suppressive miRNAs.

### 1q^AMP^ ceRNAs de-repress pro-metastatic genes by sequestering shared miRNAs

A thorough literature search identified 301 metastasis-related genes that are validated targets of the 8 tumor suppressor miRNAs sequestered by the 1q^AMP^ ceRNAs (**Figure S5A; Table S7**). Among these are 75 genes whose expression significantly correlates (FDR p<0.0001, Spearman ≥0.25) with expression of 1q^AMP^ ceRNAs in the TCGA SKCM dataset (**Figure S5B**). Moreover, 49 of the 75 putative effectors are significantly overexpressed in melanoma metastases compared to primary tumors (**Figure S5C**) and 24 of these are shared putative effectors of two or all three 1q^AMP^ ceRNAs (**Figure S5D**). Protein levels of 7 of these candidate effectors, ZEB1, ZEB2, ROCK1, ROCK2, SP1, SUZ12, and BMI1, increased upon 1q^AMP^ ceRNA overexpression in WM793 cells (**Figures 6A**). Moreover, the expression of these putative effectors was suppressed upon silencing of endogenous *CEP170, NUCKS1*, and *ZC3H11A* in 1205Lu melanoma cells (**Figure 6B** **and S5E**) and their levels were rescued when the expression of the *CEP170, NUCKS1*, or *ZC3H11A* 3’UTRs was restored (**Figure 6B**).

**Figure 6.**
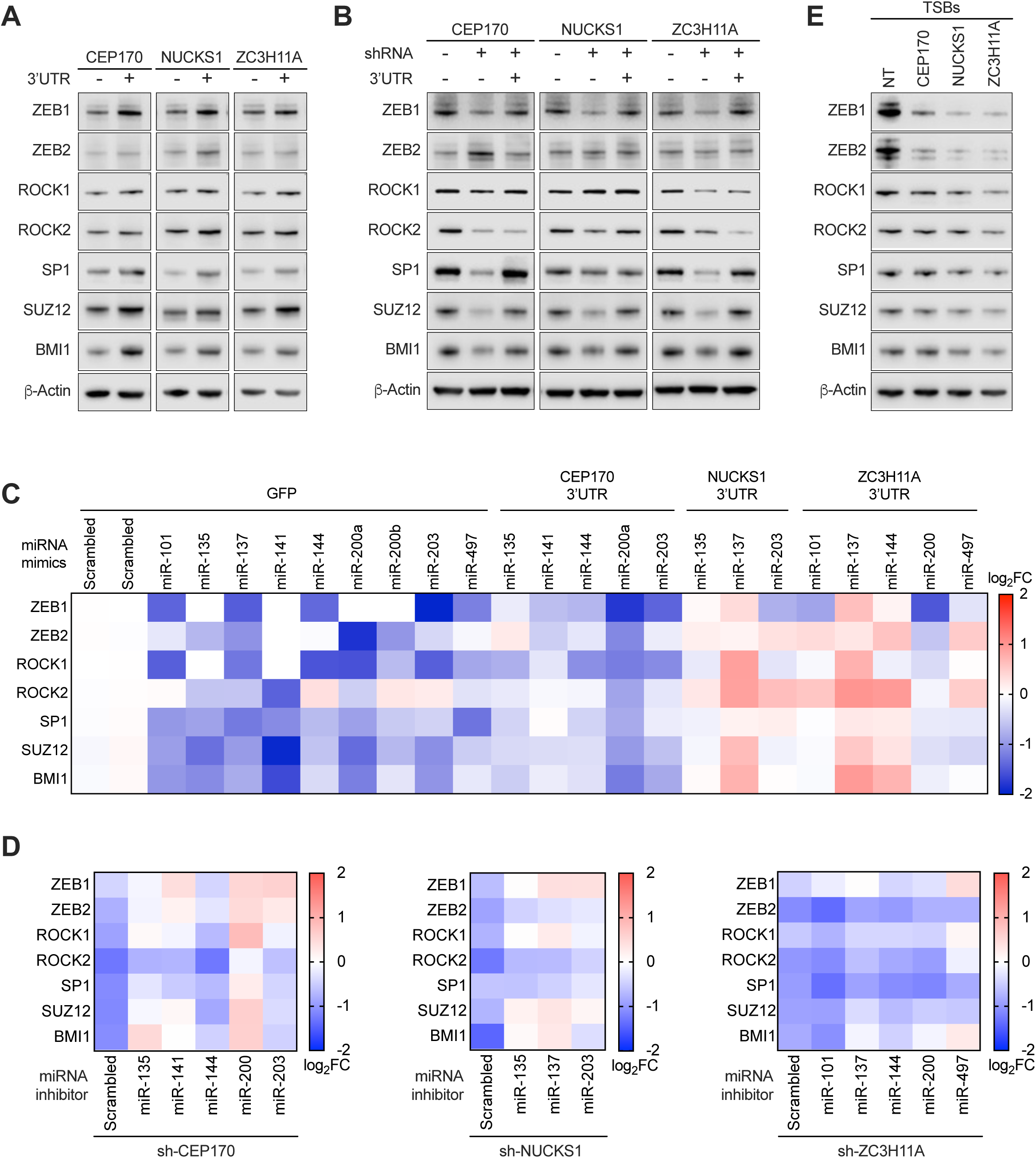
1q^AMP^ ceRNAs de-repress pro-metastatic genes by sequestering shared miRNAs. (A) Western blot of effector levels upon expression of GFP-CEP170-3’UTR, GFP-NUCKS1-3’UTR, or GFP-ZC3H11A-3’UTR in WM793 cells. (B) Western blot of effector levels upon 1q^AMP^ ceRNA silencing in WM793 cells and restoration with GFP-CEP170-3’UTR, GFP-NUCKS1-3’UTR, or GFP-ZC3H11A-3’UTR. (C) Heatmap showing expression changes of the 1q^AMP^ ceRNA effectors detected by RT-qPCR. The indicated miRNA mimics were transfected into WM793 cells expressing GFP control, GFP-CEP170-3’UTR, GFP-NUCKS1-3’UTR, or GFP-ZC3H11A-3’UTR. Color bar represents log_2_ fold change of expression. log_2_ fold change values and p-values represented in the heatmap are listed in Table S8 (D) Heatmap showing expression changes of the 1q^AMP^ ceRNA effectors detected by RT-qPCR. The indicated miRNA inhibitors were transfected into 1205Lu cells in which endogenous *CEP170, NUCKS1*, or *ZC3H11A* are knocked down. Color bar represents Log_2_ fold change of expression. Log_2_ fold change values and p-values represented in the heatmap are listed in Table S9 (E) Western blot of effector levels in1205Lu cells transfected with TSBs combinations to block the relevant MREs on endogenous *CEP170, NUCKS1*, or *ZC3H11A*. N = 3 biological replicates. See also Figure S5.

Notably, the regulation of these putative effectors by 1q^AMP^ ceRNAs was miRNA-dependent. First, transfection of WM793 melanoma cells with the respective miRNA mimics suppressed the expression of the 7 putative effectors (**Figure 6C**). Second, concomitant overexpression of the *CEP170, NUCKS1*, or *ZC3H11A* 3’UTRs limited the repression of these effectors by the miRNA mimics (**Figure 6C**). Third, transfection of 1205Lu melanoma cells with miRNA inhibitors negated the effects of silencing endogenous *CEP170, NUCKS1*, or *ZC3H11A* on the expression of these seven effectors (**Figure 6D**). Fourth, blocking MREs of endogenous *CEP170, NUCKS1*, or *ZC3H11A* mRNAs using TSBs reduced expression of these effectors (**Figures 6E** **and S5F**). Finally, individual silencing of the 7 effectors reduced invasion of 1205Lu cells (**Figure S5G**), indicating their contribution to the phenotype elicited by the 1q^AMP^ ceRNAs. Taken together, *CEP170, NUCKS1*, and *ZC3H11A* function as ceRNAs that control expression of the pro-metastatic genes *ZEB1, ZEB2, ROCK1, ROCK2, SP1, SUZ12*, and *BMI1* by sequestering shared miRNAs.

### Evidence for the 1q^AMP^ ceRNA network in multiple cancer types

Chromosome 1q copy number alterations occur in multiple other cancer types (Beroukhim et al., 2010). To investigate the contribution of the 1q^AMP^ ceRNA network to tumor progression and metastasis across cancer types we queried TCGA for concomitant copy number alterations of *CEP170, NUCKS1*, and *ZC3H11A*. Significant co-amplifications and co-gains of *CEP170, NUCKS1*, and *ZC3H11A* were identified in breast cancer, colon adenocarcinoma, hepatocellular carcinoma, lung adenocarcinoma and lung squamous cell carcinoma (**Figure 7A**). Notably, the expression of *CEP170, NUCKS1*, and *ZC3H11A* strongly correlates with the expression of *ZEB1, ZEB2, ROCK1, ROCK2, SP1, SUZ12*, and *BMI1* (**Figure 7B**), suggesting the 1q^AMP^ ceRNA network is also functional in these cancer types. Interestingly, a positive correlation between 1q^AMP^ ceRNAs and the pro-metastatic effectors *ZEB1, ZEB2, ROCK1, ROCK2, SP1, SUZ12*, and *BMI1* was also evident in some cancers lacking significant alterations of chromosome 1q copy number, including prostate adenocarcinoma, stomach adenocarcinoma, and uveal melanoma (**Figure 7B**). These findings suggest that a ceRNA network directed by the chromosome 1q *CEP170, NUCKS1*, and *ZC3H11A* ceRNAs contributes to the formation and/or progression of several cancer types.

**Figure 7.**
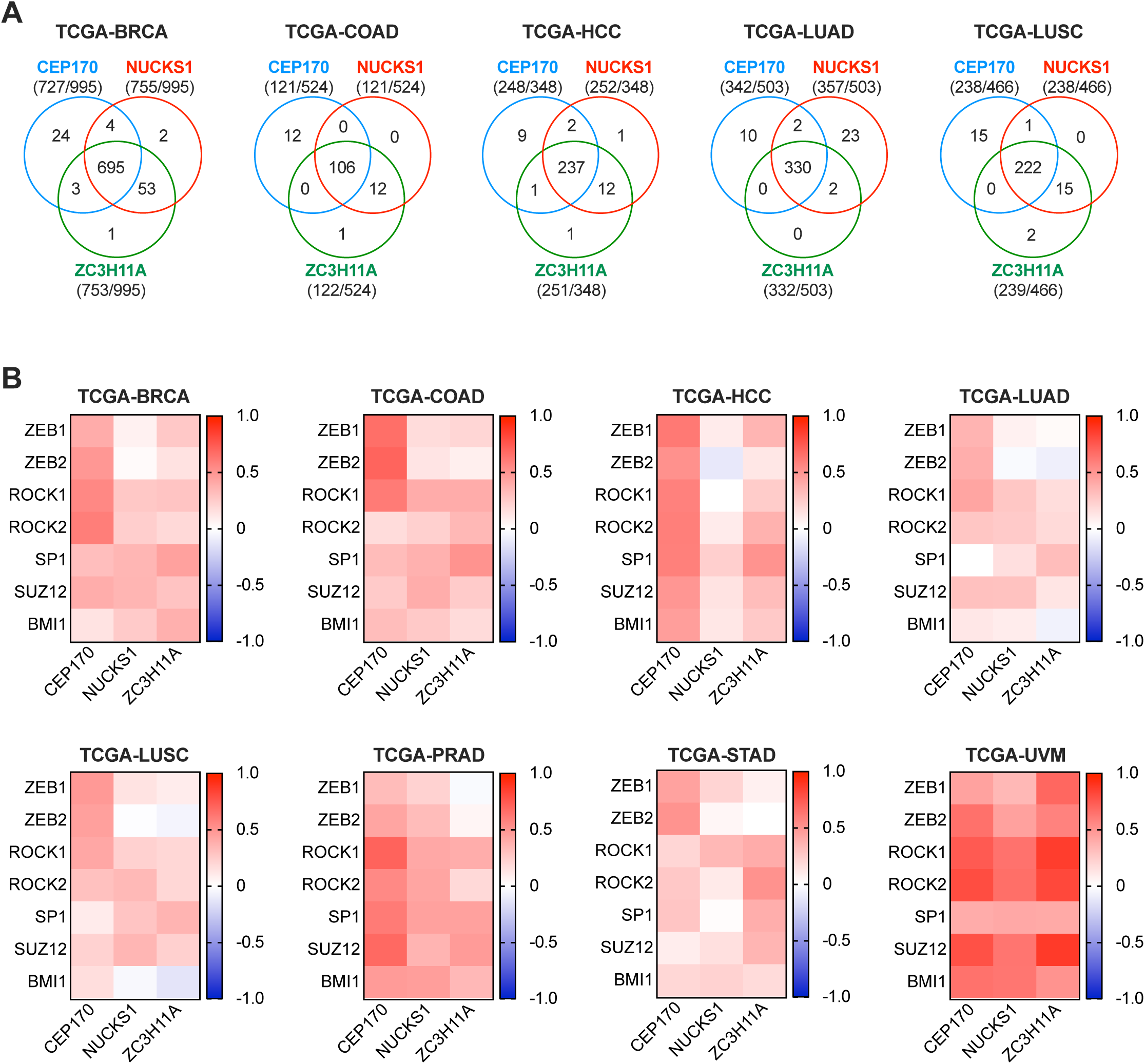
Evidence for the 1q^AMP^ ceRNA network in multiple cancer types. (A) Venn diagrams showing co-amplifications and co-gains of *CEP170, NUCKS1, ZC3H11A* in breast cancer (BRCA), colon adenocarcinoma (COAD), hepatocellular carcinoma (HCC), lung adenocarcinoma (LUAD), and lung squamous carcinoma (LUSC) in TCGA datasets. (B) Heatmaps showing the correlation between 1q^AMP^ ceRNAs and effectors in breast cancer (BRCA), colon adenocarcinoma (COAD), hepatocellular carcinoma (HCC), lung adenocarcinoma (LUAD), and lung squamous carcinoma (LUSC), stomach adenocarcinoma (STAD) and uveal melanoma (UVM). Spearman correlation values and p-values represented in the heatmap are listed in Table S10

## DISCUSSION

The studies reported herein support a paradigm where chromosome 1q copy number gains promote melanoma progression and metastasis well beyond the consequential increase in expression of certain pro-tumorigenic proteins. Instead, we show that chromosome 1q gains promote melanoma and potentially other cancer types by deregulating an oncogenic ceRNA network controlled by three mRNAs: *CEP170, NUCKS1*, and *ZC3H11A*. This oncogenic ceRNA network functions by sequestering 8 tumor suppressive miRNAs, impairing their repression of pro-metastatic target genes and thereby enabling progression to metastatic disease.

The discovery of the ceRNA concept prompted several mathematical models (Ala et al., 2013; Bosia et al., 2013; Chiu et al., 2018; Figliuzzi et al., 2013; Hausser and Zavolan, 2014; Jens and Rajewsky, 2014; Sumazin et al., 2011) and reporter-based quantitative studies (Bosson et al., 2014; Denzler et al., 2014, 2016) seeking to understand ceRNA circuits. These studies generally support the possibility of ceRNA-mediated regulation; however, some posit that a ceRNA must add a relatively large number of MREs to a network to elicit physiologically relevant changes in expression. While this contrasts with the extensive experimental evidence supporting ceRNA functions of mammalian mRNAs (Chiu et al., 2017; Gao et al., 2016; Gilot et al., 2017; Jeyapalan et al., 2011; Karreth et al., 2011; Powers et al., 2016; Rutnam and Yang, 2012; Sumazin et al., 2011; Tay et al., 2011), it suggests that specific conditions are required for a mRNA to have ceRNA activity. Thus, the prevalence of ceRNA activity among mRNAs may be moderate. Based on these considerations, we used stringent criteria - high level gains of genes having high basal expression as well as a positive correlation with putative effectors that share binding sites for at least 4 miRNAs - for our ceRNA prediction. Only 40 putative ceRNAs were identified with these criteria, indicating that only a small number of mRNAs may potently contribute as ceRNAs to the oncogenic effects of CNAs in melanoma.

ceRNAs can be classified as exclusive ceRNAs or network ceRNAs based on their mode of action. Exclusive ceRNAs are individual transcripts that sequester a single miRNA family, and unusual circumstances may be required for their activity. For instance, amplification of *MYCN* in neuroblastoma leads to >100-fold overexpression, providing almost 20% of the let-7 target site pool, and an MRE:miRNA ratio where MREs significantly exceed the number of let-7 molecules (Powers et al., 2016). Another example of an exclusive ceRNA is *TYRP1* that sequesters miR-16 in melanoma through three non-canonical target sites that do not provoke *TYRP1* mRNA decay (Gilot et al., 2017). By contrast, network ceRNAs, as highlighted by 1q^AMP^ ceRNAs, sequester multiple miRNAs that can converge on the regulation of a biological process such as metastasis. Effective miRNA sequestration can be mediated by the presence of multiple MREs and atypical expression changes or non-canonical MREs may not be strictly required. Indeed, the 1q^AMP^ ceRNAs harbor canonical 8mer and 7mer MREs for the relevant miRNAs and their expression, while more abundant than that of the sequestered miRNAs, does not reach levels observed for *MYCN*. Neither *MYCN* and *TYRP1* (Gilot et al., 2017; Powers et al., 2016) nor the 1q^AMP^ ceRNAs (data not shown) reduce miRNA levels, indicating that, given the absence of extensive target site complementarity, target RNA-directed miRNA degradation (TDMD) plays no part in the mechanism of action of these exclusive and network ceRNAs.

The network character of the 1q^AMP^ ceRNAs has two intriguing consequences. First, as *CEP170, NUCKS1*, and *ZC3H11A* share MREs for the same miRNA families, their overexpression sequesters more copies of these miRNAs. Second, two different ceRNAs can regulate the same effector through different miRNAs. Overall, this raises the ceRNA:miRNA ratio to augment miRNA sequestration. As the robustness of ceRNA crosstalk increases with the number of sequestered miRNAs (Ala et al., 2013; Chiu et al., 2018), concomitant overexpression of 1q^AMP^ ceRNAs may more potently augment the expression of their effectors. Consistent with this notion, combined overexpression of the *CEP170, NUCKS1*, and *ZC3H11A* 3’UTRs promotes migration and invasion more potently than overexpression of individual 1q^AMP^ ceRNAs. These findings support the hypothesis that the concerted deregulation of multiple ceRNAs provokes more robust effects (Smillie et al., 2018).

Chromosome 1q gains occur in various cancers in addition to melanoma. Remarkably, we found evidence for the 1q^AMP^ ceRNA network in other cancers having 1q gains and even in cancers in which 1q gains are rare. This further supports the notion that the 1q^AMP^ ceRNA network plays important pathophysiological roles. Besides the 1q ceRNA network, we identified 13 putative ceRNAs localized on chromosome 6p. While the 6p ceRNAs are predicted to regulate fewer effectors than the 1q ceRNAs, they may nevertheless cooperate to promote tumorigenesis in melanomas having 6p gains. Overall, however, ceRNA networks associated with individual CNAs may be rare, at least when our criteria are used for their prediction. Nevertheless, we foresee that additional CNA-associated ceRNA networks contributing to the development of other cancers will be identified, and their importance and oncogenic potential will be dictated by the frequency of the CNA and the expression levels of ceRNAs, miRNAs, and effectors.

In summary, we have systematically identified a complex pro-metastatic ceRNA network directly from patient level genomics data and highlight a key regulatory mechanism controlled by three genes, *CEP170, NUCKS1*, and *ZC3H11A*, that are concomitantly subjected to copy number alterations. Our study highlights how ceRNA cooperation can strongly distort regulatory networks to drive cancer progression, which we posit is a potential mechanism underlying the oncogenic effects of CNAs. Future studies of CNAs should therefore consider the contribution of ceRNA networks, especially when gains or deletions lack obvious protein-coding driver genes.

## ACKNOWLEDGMENTS

We are grateful for Karreth lab members for helpful discussions, and to J. Cleveland, A. Gomes, G. DeNicola, and Y. Tay for critical reading of the manuscript. We are grateful to A. Angarita for help with molecular cloning. This work was supported by grants from the National Institutes of Health (R03CA227349, R01CA259046) and from the Melanoma Research Alliance (https://doi.org/10.48050/pc.gr.75702) to F.A.K. This work was also supported by the Gene Targeting Core, which is funded in part by Moffitt’s Cancer Center Support Grant (P30CA076292).

## AUTHOR CONTRIBUTIONS

F.A.K conceived and supervised the project. X.X. designed and performed the majority of experiments. K.W. identified validated miRNA targets and O.V. assisted with cell line derivation. A.V., O.E., and X.Y. performed ceRNA network predictions. X.X. and F.A.K. wrote the manuscript with input from all authors.

## STAR Methods

### RESOURCE AVAILABILITY

#### Lead contact

Further information and requests for resources and reagents should be directed to and will be fulfilled by the lead contact, Florian A. Karreth.

#### Materials availability

Plasmids will be deposited at Addgene. Targeted ES cell lines/mouse strains are available upon request from the lead contact.

#### Data and code availability

- All data reported in this paper will be shared by the lead contact upon request.
- This paper does not report original code.
- Any additional information required to reanalyze the data reported in this paper is available from the lead contact upon request

### EXPERIMENTAL MODEL AND SUBJECT DETAILS

#### Cell lines and culture conditions

Human melanoma cell lines WM793 and 1205Lu were provided by M. Herlyn (Wistar). A375 were obtained from ATCC, HEK293T Lenti-X cells were purchased from Takara. Primary mouse melanoma cell lines were isolated from mouse melanomas. Briefly, tumors were washed in 70% ethanol and washed in PBS, chopped into small pieces and dissociated using the Milteny Tumor Dissociation Kit (Milteny, #130-095-929) following the manufacturer’s recommendations. Digested tissues were filtered using MACS® SmartStrainers (70 µm) (Milteny, #130-098-462) into a conical tube with 5ml of RPMI-1640 containing 5% FBS and 1% Pen/Strep to obtain a single cell suspension. Cells were collected by centrifugation, resuspended in 10 ml of RPMI-1640 containing 5% FBS and 1% Pen/Strep, and plated in a 10cm dish. After 1-2 passage, cells were transplanted subcutaneously into the flanks of NSG mice for in vivo passaging. Subcutaneous tumors from NSG mice were processed as above and used to establish the murine cell lines. All cell lines were cultured in a humidified atmosphere at 37°C and 5% CO_2_. Melanoma cell lines were cultured in RPMI-1640 (Lonza) containing 5% FBS (Sigma). HEK293T Lenti-X were cultured in DMEM (VWR) containing 10% FBS. All cell lines were confirmed to be mycoplasma-free by the MycoAlert Mycoplasma Detection Kits (Lonza).

#### Animal models

All animal experiments were conducted in accordance with an IACUC protocol approved by the University of South Florida. NSG mice were obtained from JAX (Stock No: 005557) and bred in-house. 6-week-old male and female NSG mice were randomly divided into groups (at least 5 mice per group). 1.5×10^5^ 1205Lu or 5×10^5^ WM793 melanoma cells, were intravenously injected into NSG mice via the tail vein, and lung metastasis burden was monitored after 19-22 days or 50 days, respectively, by IVIS in vivo bioluminescence imaging. 1×10^5^ A375 melanoma cells were subcutaneously injected into NSG mice, and spontaneous lung metastases were monitored after 28 days by IVIS in vivo bioluminescence imaging. For IVIS imaging, mice were anesthetized with isoflurane and then intraperitoneally injected with 100 µL Luciferin (4 mg/ml). Bioluminescent imaging was performed 10 min following Luciferin injection using the Xenogen IVIS Spectrum (Caliper Life Sciences). Mice were euthanized 3 days after IVIS imaging, and lungs were resected and fixed in Formalin. Tissue embedding in paraffin, sectioning, and H&E staining were performed by IDEXX. The quantification of lung metastasis tumor burden was performed with QuPath. ESC-GEMM chimeras were produced as previously described (Bok et al., 2019). Briefly, BP ES cells were targeted with the appropriate targeting vectors via recombination-mediated cassette exchange by the Moffitt Gene Targeting Core. Targeting was verified by PCR, and at least two positive clones were injected into Balb/c blastocysts to produce chimeras. Tumor development was induced by administering 25 mg/ml 4OHT in DMSO to the shaved back skin with a paintbrush on 2 consecutive days at 3-4 weeks of age. BP ES cells are male, but the sex of blastocysts at the time of injection is not known. Thus, either male or hermaphroditic chimeras were used. Chimeras were placed on Doxycycline-containing chow (Envigo) immediately following 4OHT administration. Chimeras were euthanized when primary tumor size reached 1 mm^3^.

### METHOD DETAILS

#### TCGA data and ceRNA prediction

Copy number and expression data from 366 Skin Cutaneous Melanoma (SKCM) metastatic cases in The Cancer Genome Atlas was analyzed. Copy number variation determined by GISTIC 2.0 (−2 = homozygous deletion; -1 = hemizygous deletion; 0 = neutral / no change; 1 = gain; 2 = high level amplification) was downloaded from cBioPortal (http://www.cbioportal.org/). Gene expression quantified from RNA-seq (TPM values generated by RSEM) was downloaded from Broad Firehose website (https://gdac.broadinstitute.org/). Genes were ranked by average TPM values across 366 metastatic cases. A total of 211 genes with high level amplification in at least 3% cases and ranked within the top 20% highest expressed genes were further considered as ceRNA candidates. Spearman correlation was calculated between candidate ceRNAs and 20,291 protein-coding genes. ceRNA-gene pairs with Spearman correlation coefficient ≥0.5 and FDR adjusted P-value <0.05 were used for miRNA binding site prediction using TargetScan (Agarwal et al., 2015). A network was constructed with gene pairs sharing at least 4 miRNAs with aggregate P_CT_ score (Friedman et al., 2009) of ≥0.2 using R package network and visualized using R package ggnet2. Connectivity of the ceRNA nodes in the network was evaluated using the Hyperlink-Induced Topic Search (HITS) algorithm implemented in R package networkR with default settings.

#### Gene set enrichment analysis

Global mRNA expression profiles of TCGA skin cutaneous melanoma dataset (TCGA-SKCM) were subject to GSEA to identify the association of *CEP170, NUCKS1*, and *ZC3H11A* with a melanoma metastasis gene signature (WINNEPENNINCKX_MELANOMA_METASTASIS_UP). For GSEA, expression of *CEP170, NUCKS1*, and *ZC3H11A* was used as phenotype, and “No_Collapse” was used for gene symbol. The metric for ranking genes in GSEA was set as ‘Pearson’, otherwise default parameters were used. GSEA was performed using GSEA 4.1.0 software.

#### Plasmid and lentivirus production

EGFP in pLenti-GFP-puro was removed with BamHI and SalI and replaced with the open reading frames (ORFs) of human *CEP170, NUCKS1, ZC3H11A*, or *AKT3*. The WPRE in pLenti-EGFP-blast, pLenti-EGFP-puro, or pLenti-EGFP-hygro was removed with SalI and EcoRI and replaced with the 3’UTRs of *CEP170, NUCKS1, ZC3H11A*, or *AKT3.* The sequences of shRNAs targeting *CEP170, NUCKS1*, or *ZC3H11A* were obtained from SplashRNA (http://splashrna.mskcc.org/) and cloned into pLenti-EGFP-miRE-puro using XhoI and EcoRI sites. To produce lentivirus supernatants, 6 µg of lentiviral vector along with 5.33 µg of Δ8.2 and 0.67 µg of VSVG helper plasmid were co-transfected into HEK293T Lenti-X cells in 10 cm dishes at 90% confluency. Lentiviral supernatant was collected 48 and 72 hours after tranfection, cleared using a 0.45 micron syringe filter, and used immediately to infect melanoma cells in the presence of 8 µg/ml polybrene or stocked in -80°C. Infected cells were selected with the appropriate antibiotics (1 µg/ml puromycin, 100 µg/ml hygromycin, or 10 µg/ml blasticidin) for 7 days.

#### Proliferation assay

800 A375 cells/well or 2,000 WM793 cells/well were plated in three 96 well plates in 100µL RPMI-1640 medium containing 5% FBS. Every other day, one plate was fized and stained cell with 50 µL crystal violet solution (0.5% crystal violet in 25% methanol) for 15 minutes. Crystal violet was discarded, and plates were gently washed twice with ddH_2_O and air dried. 150 µl of 10% acetic acid was added per well to extract crystal violet. Absorbance of extracted crystal violet was measured at 600 nM using a Glomax microplate reader.

#### Soft agar assay

1.5% and 0.8% SeaPlaque Agarose in PBS solutions were made my microwaving. 1.5% agarose was mixed at a 1:1 ratio with RPMI-1640 medium to achieve 0.75% agarose, followed by plating 1 ml per well in a 12 well plate and letting agarose solidify at room temperature. Melanoma cells were trypsinized, counted and 500 cells were resuspended in 500 µl of RMPI-1640 medium containing 20% FBS. Cells were then mixed with 500 µl of 0.8% agarose (0.4% agarose and 10% FBS final concentrations) and plated atop the solidified 0.75% agar. Top layer was allowed to solidify at room temperature, followed by culture at 37°C with 100 µl of fresh medium added every 3 days. Pictures are taken after 10-15 days and analyzed with Image J.

#### Transwell migration and invasion assay

To pre-coated transwells with Matrigel, Matrigel was thawed on ice for at least 2 hours, diluted to 5% with ice-cold basic RPMI-1640 medium, and gently added to transwell inserts (50 µl/insert) and solidified at 37°C for 30 min. Melanoma cells were trypsinized and resuspend in RPMI-1640 medium. For transwell invasion assays, 5,000 A375 cells, 15,000 1205Lu cells, or 30,000 WM793 cells were plated per Matrigel-coated insert in 200 µl of RPMI-1640 without FBS. For transwell migration assay, 3,000 A375 cells, 10,000 1205Lu cells, or 20,000 WM793 cells were plated per insert in 200 µl of RPMI-1640 without FBS. 500 µl RPMI-1640 medium containing 15% FBS were added into the bottom well. Plates were incubated at 37°C in a humidified incubator for 48 hr. Media was discarded, inserts were gently washed once with PBS, and cells were fixed with 500 µl fixing solution (ethanol:acetic acid = 3:1) in the bottom well outside the insert for 10 minutes. Inserts were washed once with PBS, and cells were stained in 500µL staining solution (0.5% crystal violet) for 30 minutes. Inserts were washed with tap water twice and non-migrated cells or Matrigel on the top side of the insert were carefully removed with cotton swabs, and inserts were air dried overnight. Pictures were taken at 200x magnification and cell numbers quantified.

#### miRNA reporter assay

Reverse complimentary sequences of candidate miRNAs (Table S2) were cloned into psiCHECK2 dual-luciferase reporter vector at the XhoI and NotI sites downstream of Renilla luciferase. 2×10^4^ cells/well were seeded in 96-well plates and allowed to settle for 24 hours. For each well of 96 well plate, 15 ng reporter plasmid were transfected using 0.5 µl Jetprime in 5 µL Jetprime buffer. 48 hr after transfection, cells were washed with PBS and lysed in 50 µl of lysis buffer (1x PLB, made from 5x PLB contained in the Dual Luciferase Reporter Assay System kit). Plates were rocked at room temperature for 15 min, and 20 µl of cell lysates were transferred to white-walled 96-well plates. Wash the injectors of the Glomax plate reader twice with ddH_2_O, prime Injector 1 with buffer LAR II (for Firefly Luciferase) and prime Injector 2 with buffer STOP’n’GLO (for Renilla Luciferase, both contained in the Dual Luciferase Reporter Assay System kit). Set up the procedure for each well (Injector 1 volume: 50 µl, waiting duration: 2 seconds, reading integration: 5 seconds, Injector 2 volume: 50 µl, waiting duration: 2 seconds, reading integration: 5 seconds) then read the plate well by well in the luminometer automatically. Empty psiCHECK-2 control lacking MREs and appropriate vector or scramble controls were included in the experimental setup. Renilla luciferase values were normalized to Firefly luciferase value to control for transfection efficiency. To analyze data, normalized Luciferase values of the MRE-bearing reporter were first compared to the empty reporter in vector or scramble control cells, and then the normalized Luciferase values of MRE-bearing reporter were compared to ceRNA overexpression or knockdown conditions.

#### RNA pulldown

Biotinylated oligonucleotide probes targeting GFP were designed using the online tool (https://www.biosearchtech.com/support/tools/design-software/chirp-probe-designer) and synthesized by Eton Bioscience. 1×10^7^ cells were crosslinked using the UV cross-linker (Spectrolinker) with 600-1200J/cm^2^. Cells were washed with PBS, collected in 1.5 ml tube, and lysed with 500-800 µl lysis buffer (TBS (pH7.4-7.6), 1 % Triton-X100, 0.1-0.5 % deoxycholate, 0.1-0.5 % SDS) containing 1% protease inhibitor cocktail and 0.1-0.5 % RNase inhibitor on ice for 15 min with repeated vortexing every 5 min. Lysates were cleared by centrifugation at 12,000g for 5 min at 4°C and transferred to new 1.5ml tubes. 100-200 µl of cell lysate were diluted 1:2 in hybridization buffer (500mM NaCl, 1% SDS, 100mM Tris (pH7.0), 10mM EDTA, 15% formamide). 1-2 nmol of probe mixture was added and mixed by end-to-end rotation at 37°C for 4 hours. Streptavidin beads were pre-cleared twice by adding 1 ml of hybridization buffer and collected using a magnetic rack. 20µl of Streptavidin beads were added per sample, mixed by end-to-end rotation at 37°C for 30 minutes, and washed 5 times in wash buffer for 5 minutes each at room temperature. Streptavidin beads were resuspended in 100 µl Proteinase K buffer (100mM Tris pH7.5, 200mM NaCl, 2mM EDTA, 1% SDS) containing Proteinase K (1 mg/ml), incubated at 55°C for 30 minutes, and boiled for 10 min. RNA was then isolated with the Qiagen miRNeasy kit.

#### miRNA mimics, miRNA inhibitors, Target Site Blockers (TSB)

miRNA mimics and miRNA inhibitors were purchased from Horizon Discovery and resuspended in ddH_2_O to prepare 20µM solutions. miRNA mimics or miRNA inhibitors at 10 nM or 20 nM final concentration, respectively, were transfected into melanoma cells using JetPrime according to the manufacturer’s recommendations. 48 hr after transfection, transwell invasion assays were performed or RNA was isolated.

Target Site Blockers: TSBs targeting the MREs of (i) miR-135, miR-141/200a, miR-144, or miR-203 on the *CEP170* 3’UTR; (ii) miR-135, miR-137, or miR-203 on the *NUCKS1* 3’UTR; and (iii) miR-101/137/144, or miR-200b/200c on the *ZC3H11A* 3’UTR were obtained from Qiagen. TSBs were resuspended in ddH_2_O to prepare a 50 µM solution. TSBs were transfected into melanoma cells at a final concentration of 50nM. For xenograft metastasis assays, melanoma cells were intravenously injected into NSG mice 24 hr after TSB transfection. Transwell invasion assays were performed 48 hr after TSB transfection. RNA and protein were isolated 72 hr and 96 hr, respectively, after TSB transfection.

#### Droplet digital PCR (ddPCR)

RNA was isolated from 1×10^6^ cells using Trizol using the manufacturer’s recommendation. RNA was reverse transcribed into cDNA using the TaqMan MicroRNA Reverse Transcription Kit (Thermo Fisher Scientific) (miRNA) or PrimeScrip RT Master Mix (Takara) (mRNA). cDNA samples (10fg∼100ng/reaction), Taqman probes (FAM, 20x), ddPCR supermix (2x) and ddH_2_O were combined in a total volume of 22 µl in 96-well plates, and plates were sealed, centrifuged for 30 sec at 1,000rpm, and loaded into the Generator along with required consumables (DG32 cartridges, tips, waste bins, droplet plate, and oil) to partition the samples into droplets. Plates the containing droplets were sealed with foil, and PCR was performed to end point (∼40 cycles) using the conditions outlined in Table S11. Plates were then loaded into the QX200 Reader to quantify positive and negative droplets in each sample and to plot fluorescence signal droplet by droplet. Results are visualized in Quantasoft Software. The fraction of positive droplets in a sample determines the concentration of target in copies/µl.

#### RT-qPCR

RNA was extracted from cells using TRIzol (Invitrogen) following protocols supplied by the manufacturer. First-strand cDNA was generated with PrimeScript RT Master Mix (Takara). qPCR was performed on StepOnePlus™ Real-Time PCR System (Thermo Fisher), and PerfeCTa SYBR green Fastmix (Quantabio) was used for gene detection. The sequences of primers were obtained from PrimerBank (https://pga.mgh.harvard.edu/primerbank/index.html) and are listed in Table S2.

#### Western blot

15 µg of protein were loaded onto NuPAGE 4%-12% precast gels and separated by electrophoresis with NuPAGE running buffer at constant 120V for approximately 2 hr. Proteins were transferred to nitrocellulose membranes with transfer buffer at constant 300mA for 2 hr. Membranes were blocked in 5% non-fat dry milk solution in TBST for 30 min and then incubated with one of the following primary antibodies overnight at 4°C: anti-ZEB1 (1:1,000), anti-ZEB2 (1:1,000), anti-ROCK1 (1:1,000), anti-ROCK2 (1:1,000), anti-SP1 (1:1,000), anti-SUZ12 (1:1,000), or anti-BMI1 (1:1,000). Anti-beta-actin (1:3,000) was blotted as a loading control. Membranes were washed 3 times with TBST for 10 min, followed by incubation with HRP-conjugated secondary antibodies (1:2,000) for 1 hr room temperature. After three washes in TBST, chemiluminescence substrate (1:1) was applied to the blot for 4 min and chemiluminescence signal was captured using a LI-COR imaging system.

#### Quantification and statistical analysis

Proliferation assays are presented as mean ± SD. For soft agar, transwell migration, and invasion assays, 3 or 4 random fields were quantified for each well and data are presented as mean ± SEM. For luciferase reporter assays and RNA pulldown assays data are presented as mean ± SEM. Droplet digital PCR data were generated using 4 replicates and presented as mean ± SD. All experiments were performed at least 3 times with 3-4 technical replicates. Statistical analyses were performed by Student’s unpaired t test. Data of in vivo imagining of lung metastasis are presented as mean ± SEM. Survival analysis of ESC-GEMMs was based on a cohort of 58 mice (22 TRE-GFP mice, 10 TRE-CEP170 mice, 11 TRE-NUCKS1 mice, and 15 TRE-ZC3H11A mice) and the p values were calculated based on a log-rank test. A p value of < 0.05 was considered statistically significant.

## SUPPLEMENTAL TABLES

**Supplemental Table S1:** List of 211 candidate ceRNAs

**Supplemental Table S2:** List of miRNA families predicted for the candidate ceRNAs

**Supplemental Table S3:** List of ceRNA/effector gene pairs

**Supplemental Table S4:** HITS analysis

**Supplemental Table S5:** Expression of ceRNAs in melanocyte and melanoma cell lines

**Supplemental Table S6:** Luc-MRE reporter assay results for sponging of predicted miRNAs

**Supplemental Table S7:** List of validated pro-metastatic targets of sponged miRNAs

**Supplemental Table S8:** Log_2_ fold change of expression of effectors upon ceRNA 3’UTR overexpression ± miRNA mimics

**Supplemental Table S9:** Log_2_ fold change of expression of effectors upon ceRNA silencing ± miRNA inhibitors

**Supplemental Table S10:** Spearman correlations of 1q^AMP^ ceRNAs and effectors across 8 cancer types

**Supplemental Table S11:** RT-qPCR primer sequences, MRE reporter sequences, transfection conditions, ddPCR procedure

## SUPPLEMENTAL FIGURE LEGENDS

**Figure S1.**
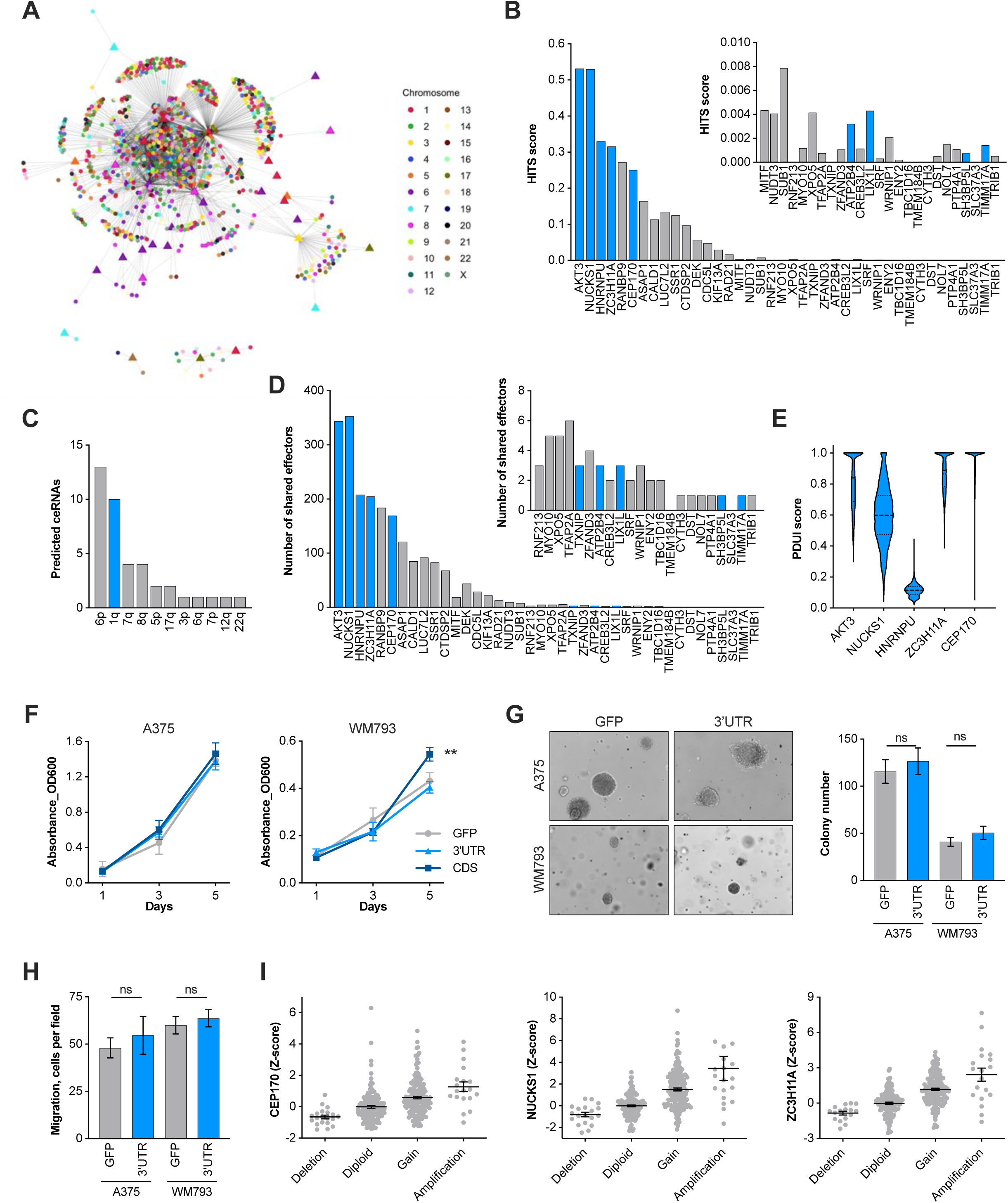
CEP170, NUCKS1, and ZC3H11A, but not AKT3 and HNRNPU, are potential ceRNAs on chromosome 1q, related to Figure 1. (A) The metastatic melanoma ceRNA network. Central triangles represent gained/amplified ceRNA genes and peripheral circles represent putative effectors. (B) HITS score of the 40 predicted ceRNAs in melanoma. The insert graph shows a close-up of the 25 bottom-ranked ceRNA genes. Predicted ceRNAs encoded by genes localized on chromosome 1q are highlighted in blue. (C) The number of predicted ceRNA genes by genomic localization. (D) The number of shared effectors of the 40 predicted ceRNAs in melanoma. The insert graph shows a close-up of the 22 bottom-ranked ceRNA genes. Predicted ceRNAs encoded by genes localized on chromosome 1q are highlighted in blue. (E) Percentage Distal Usage Index (PDUI) scores of *AKT3, NUCKS1, HNRNPU, ZC3H11A*, and *CEP170*, the top ranked ceRNAs localized on chromosome 1q. (F) Proliferation assay of A375 and WM793 cells overexpressing GFP-AKT3-3’UTR, the AKT3 coding sequence (CDS) or a GFP control. (G) Anchorage-independent growth in soft agar of A375 and WM793 cells overexpressing the GFP-AKT3-3’UTR or a GFP control. (H) Migration in a transwell assay of A375 and WM793 cells overexpressing the GFP-AKT3-3’UTR or a GFP control. (I) Correlation between genomic alteration and gene expression of *CEP170, NUCKS1*, and *ZC3H11A*. In (F-H), N = 3 biological replicates; two-sided t test. Values represent mean ± SEM. ** p < 0.01, ns not significant.

**Figure S2.**
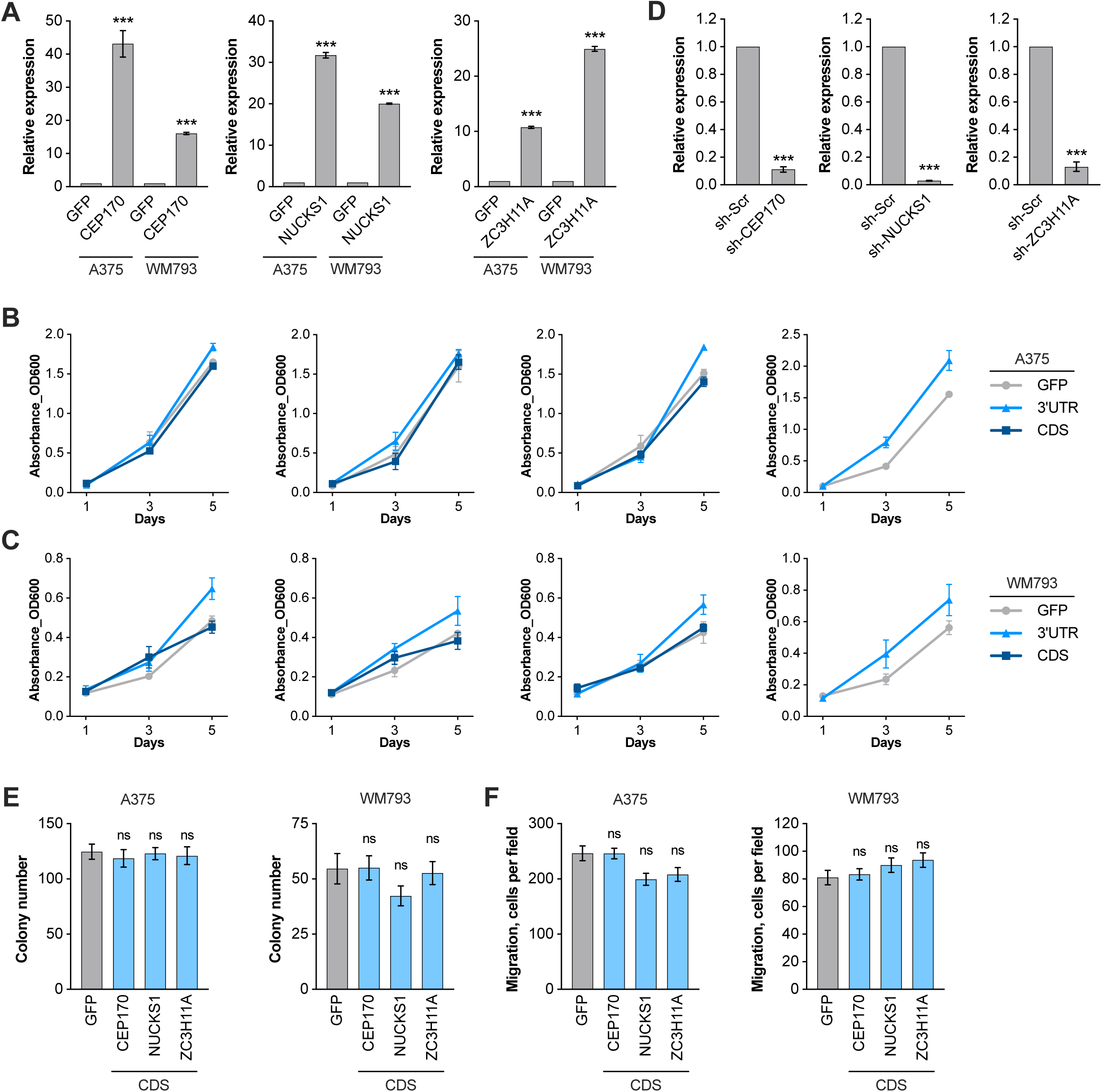
Quantification of 1q^AMP^ ceRNA overexpression and silencing, and effect of 1q^AMP^ ceRNA CDS on melanoma cell biology, related to Figure 2. (A) RT-qPCR showing the relative levels of the 1q^AMP^ ceRNA 3’UTRs in WM793 and A375 cells expressing GFP-CEP170-3’UTR, GFP-NUCKS1-3’UTR, or GFP-ZC3H11A-3’UTR. Expression was normalized to the GFP control cells. (B) Proliferation assays of A375 cells expressing GFP, the 1q^AMP^ ceRNA 3’UTRs (GFP-CEP170-3’UTR, GFP-NUCKS1-3’UTR, or GFP-ZC3H11A-3’UTR), or the 1q^AMP^ ceRNA CDS. (C) Proliferation assays of WM793 cells expressing GFP, the 1q^AMP^ ceRNA 3’UTRs (GFP-CEP170-3’UTR, GFP-NUCKS1-3’UTR, or GFP-ZC3H11A-3’UTR), or the 1q^AMP^ ceRNA CDS. (D) RT-qPCR showing the knock down efficiency of *CEP170, NUCKS1*, or *ZC3H11A* in 1205Lu cells. (E) Quantification of anchorage-independent growth in soft agar of A375 and WM793 cells overexpressing the 1q^AMP^ ceRNA CDS. (F) Quantification of transwell migration assay of A375 and WM793 cells overexpressing the 1q^AMP^ ceRNA CDS. N = 3 biological replicates; two-sided t test. Values represent mean ± SEM. * p < 0.05, ** p < 0.01, *** p < 0.001, ns not significant.

**Figure S3.**
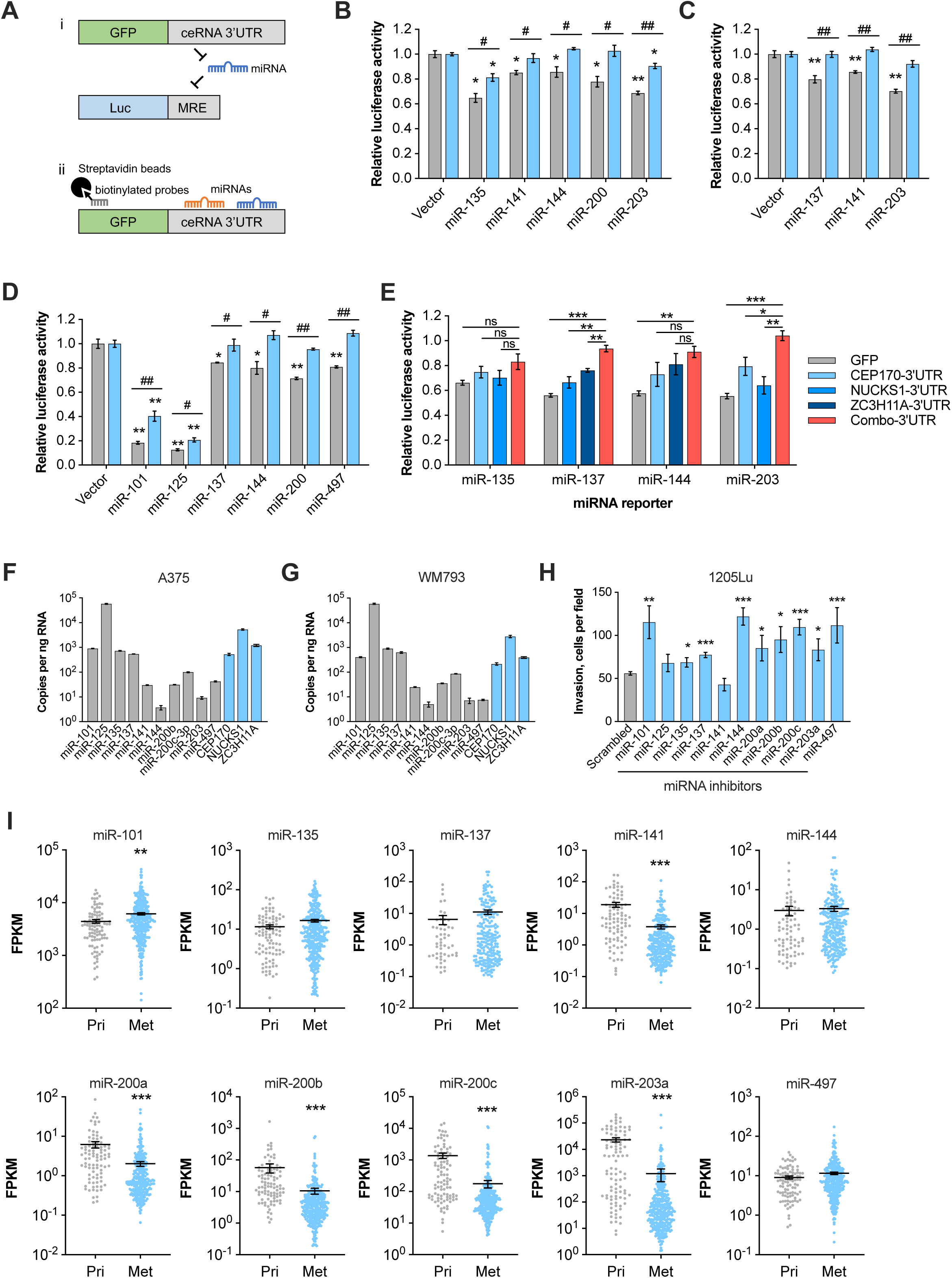
Identification of tumor suppressive miRNAs that are sequestered by the 1q^AMP^ ceRNAs, related to Figure 4. (A) (i)Outline of the MRE luciferase reporter assay. MREs were cloned downstream of Renilla luciferase, leading to repression by endogenous miRNAs. Co-expression of 1q^AMP^ ceRNAs will result in de-repression of the reporter constructs. (ii) Outline of the RNA pulldown assay. Biotinylated GFP probes were used to pull down GFP, GFP-CEP170-3’UTR, GFP-NUCKS1-3’UTR, and GFP-ZC3H11A-3’UTR with Streptavidin beads and bound miRNAs were detected by RT-qPCR. (B) Luciferase assays using dual luciferase reporters harboring MREs for miR-135-5p, miR-141-3p, miR-144-3p, miR-200-3p, or miR-203-3p in A375 cells expressing GFP or GFP-CEP170-3’UTR. Luciferase activity was normalized to the reporter without MRE (vector). (C) Luciferase assays using dual luciferase reporters harboring MREs for miR-135-5p, miR-137, or miR-203-3p in A375 cells expressing GFP or GFP-NUCKS1-3’UTR. Luciferase activity was normalized to the reporter without MRE (vector). (D) Luciferase assays using dual luciferase reporters harboring MREs for miR-101-3p, miR-125-5p, miR-137, miR-144-3p, miR-200-3p, or miR-497-5p in A375 cells expressing GFP or GFP-ZC3H11A-3’UTR. Luciferase activity was normalized to the reporter without MRE (vector). In (B-D), asterisks (*, **) indicate comparison of Luc-MRE reporter vs. empty control Luc reporter (Vector) and pound symbols (^#^, ^##^) indicate comparison of 3’UTR vs. GFP. (E) Luciferase assays using dual luciferase reporters harboring MREs for miR-135-5p, miR-137, miR-144-3p, or miR-200-3p in WM793 cells expressing the 1q^AMP^ ceRNA 3’UTRs individually or in combination. Luciferase activity was normalized to the reporter without MRE (vector), not shown. (F-G) Copies of the indicated mature miRNAs and endogenous *CEP170, NUCKS1, ZC3H11A* per ng RNA were quantified by Droplet Digital PCR in A375 (F) and WM793 (G) cells. (H) Quantification of transwell invasion assays with 1205Lu cells transfected with the indicated miRNA inhibitors. (I) Expression of the indicated miRNAs in primary and metastatic melanoma samples in the TCGA melanoma dataset. In (B-D, H), N = 3 biological replicates. Two-sided t test; values represent mean ± SEM. * p < 0.05, ** p < 0.01, *** p < 0.001, ^#^ p < 0.05, ^##^ p < 0.01.

**Figure S4.**
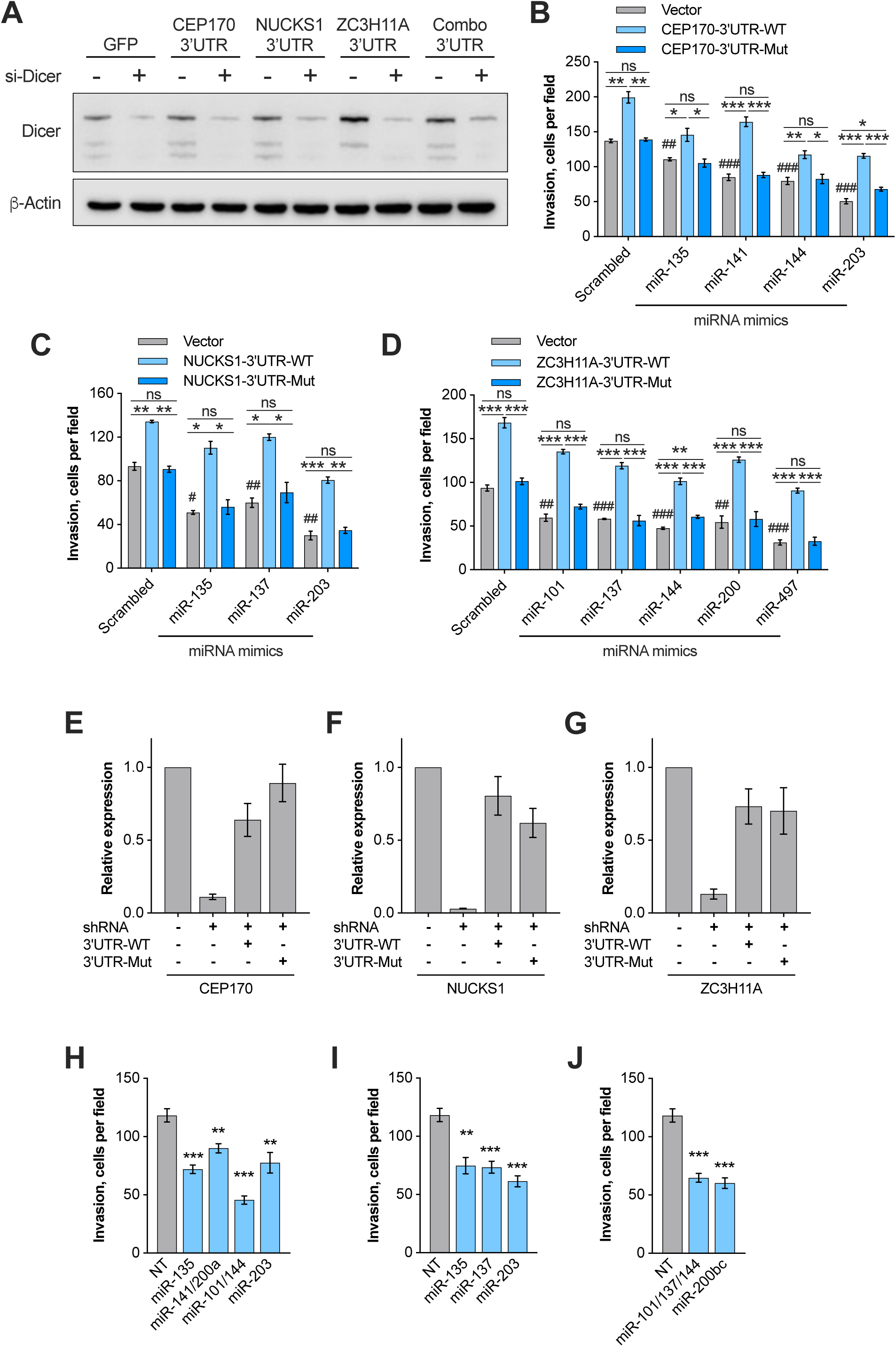
The oncogenic effect of the 1q^AMP^ ceRNAs depends on miRNA binding, related to Figure 5. (A) Western blot validates the siRNA-mediated silencing of Dicer in 1205Lu cells expressing 1q^AMP^ ceRNAs. (B-D) Quantification of transwell invasion of A375 cells upon transfection with the indicated miRNA mimics. A375 cells were engineered to overexpress GFP, or wildtype or MRE-mutant GFP-CEP170-3’UTR (B), GFP-NUCKS1-3’UTR (C), or GFP-ZC3H11A-3’UTR (D). Asterisks indicate comparisons of GFP vs. 3’UTR-WT vs. 3’UTR-Mut and pound symbols indicate comparisons of miRNA mimics vs. scrambled control in GFP control samples (blue bars). (E-G) RT-qPCR to assess restoration of 1q^AMP^ ceRNA expression upon wildtype or MRE-mutant GFP-CEP170-3’UTR (E), GFP-NUCKS1-3’UTR (F), or GFP-ZC3H11A-3’UTR (G) expression in 1205Lu cells having silenced endogenous *CEP170, NUCKS1*, or *ZC3H11A*, respectively. Asterisks indicate comparisons of 3’UTR expression in 3’UTR-WT/Mut restored vs. ceRNA shRNA and pound symbols indicate comparison of ceRNA shRNA vs. shRNA control. (H-J) Quantification of transwell invasion of 1205Lu cells transfected with the indicated target site blockers (TSBs) to block MREs on endogenous CEP170 (H), NUCKS1 (I), or ZC3H11A (J). Asterisks indicate comparisons of TSBs (light blue bars) vs. control TSB (gray bars). N = 3 biological replicates; two-sided t test. Values represent mean ± SEM. * p < 0.05, ** p < 0.01, *** p < 0.001, ^#^ p < 0.05, ^##^ p < 0.01, ^###^ p < 0.001, ns not significant.

**Figure S5.**
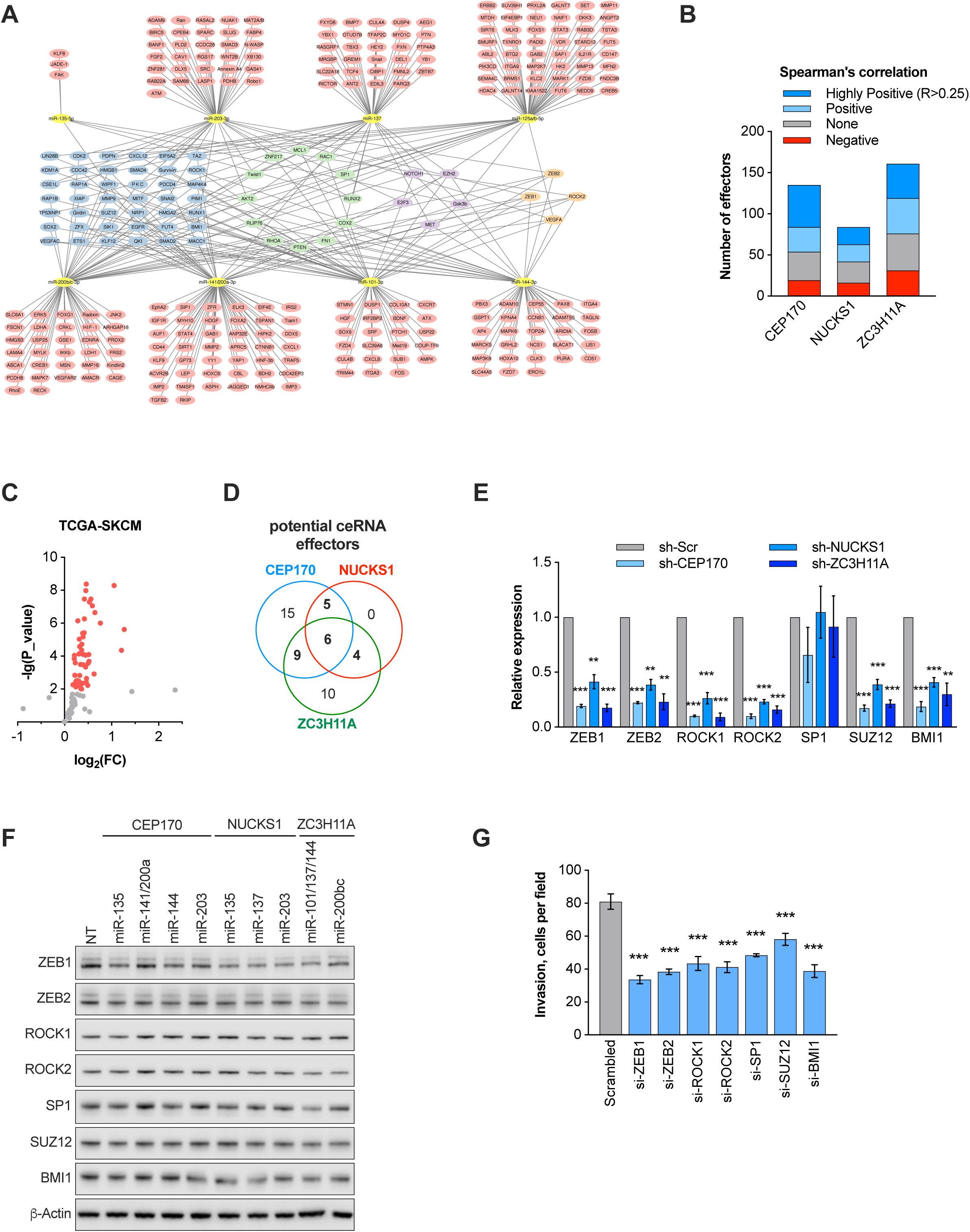
The 1q^AMP^ ceRNAs regulate expression of pro-metastatic effectors, related to Figure 6. (A) 301 metastasis associated genes are validated targets of one or more of the miRNAs sequestered by the 1q^AMP^ ceRNAs, based on published literature. (B) The expression correlation between the 301 miRNA targets and *CEP170, NUCKS1*, or ZC3H11A in the TCGA cutaneous melanoma dataset are shown. 72 genes are highly correlated (R>0.25) and are potential ceRNA effectors. (C) Volcano plot showing the differential expression of potential ceRNA effectors in primary and metastatic melanoma samples in TCGA dataset. 49 potential effectors are significantly higher expressed in metastatic melanoma compared to primary melanoma. (D) Venn diagram showing that expression of 24 out of 49 putative effector correlates with two or three of the 1q^AMP^ ceRNAs. (E) RT-qPCR showing expression of the indicated effectors in 1205Lu melanoma cells upon silencing endogenous *CEP170, NUCKS1*, or *ZC3H11A*. Effector expression was normalized to cells expressing a scrambled control shRNA (sh-Scr). (F) Western blot showing the protein expression of the indicated effectors in 1205Lu melanoma cells following treatment with individual TSBs. (G) Quantification of invasion of 1205Lu cells upon silencing of the indicated effectors with siRNAs. Values represent mean ± SEM. *** p < 0.001.

